# Enhancing viscosity control in antibody formulations: A framework for the biophysical screening of mutations targeting solvent-accessible hydrophobic and electrostatic patches

**DOI:** 10.1101/2024.03.07.583917

**Authors:** Georgina B Armstrong, Vidhi Shah, Paula Sanches, Mitul Patel, Ricky Casey, Craig J Jamieson, Glenn A Burley, William J Lewis, Zahra Rattray

## Abstract

The formulation of high-concentration monoclonal antibody (mAb) solutions in low dose volumes for autoinjector devices poses challenges in manufacturability and patient administration due to elevated solution viscosity. In the current study, we present a systematic experimental framework for the computational screening of molecular descriptors to guide the design of mutants with modified viscosity profiles accompanied by experimental evaluation. Our observations using a model anti-IL8 antibody reveal that the reduction in viscosity is influenced by the location of hydrophobic interactions, while targeting positively charged patches in mAb1 leads to the most significant viscosity increase compared to the wild-type mAb. We conclude that existing *in silico* predictions of physicochemical properties exhibit poor correlation with experimental parameters for antibodies with suboptimal developability characteristics, emphasizing the necessity for comprehensive case-by-case evaluations of mAbs. This approach aids in the rational design of mAbs with tailored solution viscosities, ensuring improved manufacturability and patient convenience in self-administration scenarios.

## Introduction

In the realm of modern medicine, therapeutic monoclonal antibodies (mAbs) have emerged as indispensable tools in the arsenal against chronic diseases such as diabetes, cancer, and autoimmune disorders. Empowering patients with self-administration regimens, subcutaneous injection has become the cornerstone of delivering these life-changing therapies, necessitating formulation design strategies to accommodate small injection volumes. However, this pursuit of patient convenience presents a formidable challenge: how to achieve high mAb formulation concentrations (>100 mg/mL) without succumbing to the challenges of elevated aggregation propensity and solution viscosity.

The viscosity of mAb formulations, a critical parameter governing dosing and delivery efficacy, is intricately linked to protein-protein interactions arising from the mAb amino acid sequence and formulation excipient composition.

High concentrations exacerbate these interactions, leading to increased aggregation risk and elevated formulation viscosity (>30 centipoise).^1^ High mAb formulation concentrations result in an exponential increase in protein-protein interactions leading to a higher propensity for aggregation. Diffusion interaction coefficients (k_D_) are used to measure protein-protein interactions and colloid stability, with high viscosity mAbs generally exhibiting large negative k_D_ values (attractive).^2–4^

In this pursuit, various strategies have been employed to modulate protein-protein interactions and mitigate mAb solution viscosity. These strategies have ranged from the alteration of electrostatic properties by changing formulation buffer pH and salt composition, to employing viscosity reducing excipients (e.g., amino acids) to increase the solubility of partially folded and unfolded states.^5–7^ Furthermore, advancements in sequence-based engineering offer a promising avenue for targeting solvent-accessible electrostatic patches on the mAb surface, with the potential to revolutionize the mAb design landscape.

In the emerging area of precision medicine, the integration of high throughput *in silico* predictions and molecular triaging approaches holds immense potential in streamlining early-stage discovery campaigns.^8–10^ By elucidating the intricate relationship between mAb molecular descriptors and developability risks,^11^ these cutting-edge approaches empower researchers to more expediently identify candidate mAbs with superior physicochemical properties, paving the way for more agile drug development pipelines with less attrition.

Current landscape analyses and models defining optimal developability for mAbs are based on clinically approved drug products with optimal characteristics. However, amidst these advancements, it is imperative to broaden our focus beyond clinically approved mAbs and encompass those with unknown or sub-optimal developability characteristics. In doing so, we expand our understanding of how to navigate high formulation concentration solution viscosity more effectively, ultimately enhancing the success rate of mAb drug development endeavors.^2^

In this manuscript, we harness a range of computational molecular descriptors as a guiding tool for the design and triage of a mutant mAb panel disrupting solvent-accessible hydrophobic and electrostatic surface patches. Through a combined computational and experimental framework, we examine the relationship between single-point mutations and the biophysical properties of a model antibody, mAb1. Our findings show site-specific and strategy-dependent impact of mutations based on surface patch composition, offering an insight into downstream effects of molecular alterations. Here, we report significant alterations in surface potential from single-point mutations in the variable region and observe favourable developability characteristics for hydrophobic or negative patch-disrupting mutants compared to the mAb1 wild-type. We observe correlations between hydrophobicity-based molecular descriptors as well as colloidal parameters in predicting hydrophobicity-driven self-associations, impacting solution viscosity at high mAb concentrations.

## Results

### Generation of the mAb1 mutant panel

Using homology models of mAb1, we compared the impact of targeting solvent-accessible hydrophobic and charged patches on solution viscosity at high mAb concentration.^12–14^ Patch analysis of a mAb1 wild-type (WT) IgG1 homology model identified residues contributing to positive, negative, and hydrophobic patches as potential candidates for mutation. We then determined physicochemical molecular descriptors and conducted patch analyses on all mutants.

### Homology modelling and patch analysis of WT mAb1

We constructed homology models of the full mAb1 structure and the Fv fragment of WT mAb1 in the MOE molecular modelling suite.^15^ Since the Fab crystal structure (PDB 5OB5) matched the framework and CDR regions perfectly, patch analysis was conducted on resulting structures (**Figure 1A**). The electrostatic potential mapped onto the mAb1 surface (**Figure 1B**), shows negative, positive, and hydrophobic patch distributions.

**Figure 1.**
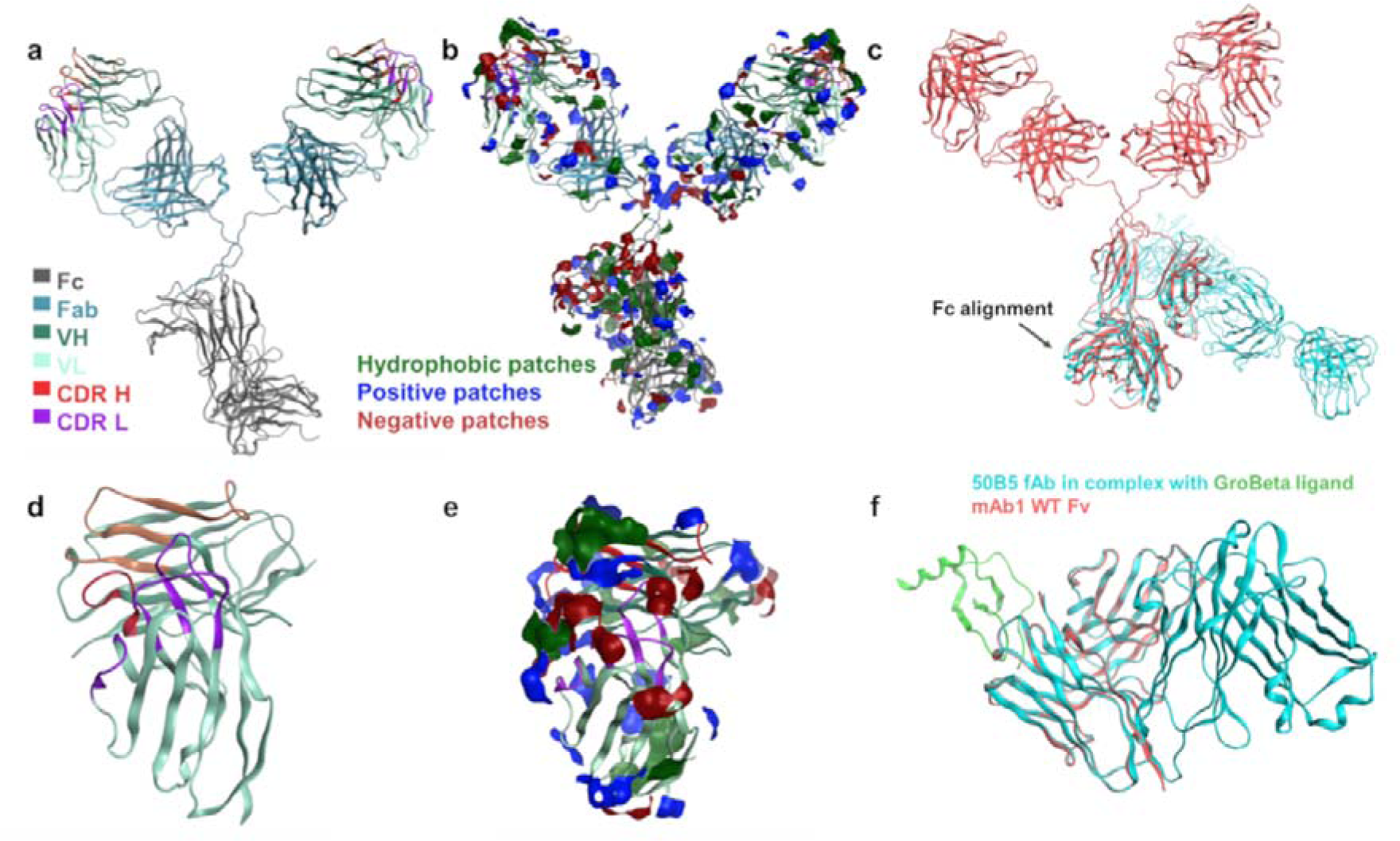
Homology models of mAb1. **a,** the full IgG structure was modelled using the PDB 5OB5 template for the Fab region and IgG model in the MOE platform The Fc (grey), constant light chain 1 and heavy chain 1 (blue), variable heavy chain (dark green) variable light chain (light green), heavy chain CDRs (red) and light chain CDRs (purple) were labelled using Kabat annotation. **b,** the hydrophobic (green), positive (blue), and negative (red) patches applied onto the full IgG1 homology model to demonstrate the exposed surface charges and accessible non-polar regions as potential sites for promoting protein-protein interactions. **c,** superimposition, and alignment of the mAb1 WT full IgG homology model (pink) onto the template 1IGY PDB IgG1 structure (blue) to model the Fc structure. **d,** the Fv region was modelled separately and used for most molecular descriptor calculations. **e,** patch analysis of the Fv to aid identification of candidate sites for single-point mutation. **f,** superimposition and alignment of the mAb1 WT Fv homology construct (pink) onto the template 5OB5 PDB fAb structure (blue) that was in complex with the GroBeta ligand (green).

Overall, we observed the largest contribution to the surface potential of WT mAb1 IgG (**Figure *1B***) from hydrophobic (3,790 Å^2^), positive (2,940 Å^2^) and negative (2,250 Å^2^) patches, with a net charge of +22.68 C (pH 6). A similar trend was seen with the mAb1 Fv (**Figure 1C and D**), with surface area coverage of 520, 160, and 50 Å^2^ for hydrophobic, positive and negative patches, respectively, and a net charge of +0.05 C (pH 6). We then identified mutant residues in the mAb1 framework and CDRs that would significantly disrupt hydrophobic, positive, and negative patches (**Supplementary Table S1**), potentially influencing protein-protein interactions and self-association.

### Patch analysis of mAb1 mutants

We explored the effects of single point mutations on the mAb1 charge and hydrophobic patch distributions, by introducing Fv point mutations. Employing strategies targeting positive, hydrophobic, and negative patches, we observed changes in electrostatic surface potentials following framework region and CDR mutations (**Figure 2**).^12–14^ The mAb1 Fv carries a net positive charge (+0.05 C, pH 6.0), with heterogeneous surface charge distribution, resulting in asymmetry between heavy and light chain net charges (3.93 C and - 1.23 C, respectively). Patch analysis of the WT Fv revealed significant hydrophobicity (520 Å^2^) with prevalent surface coverage by positive patches (blue).

**Figure 2.**
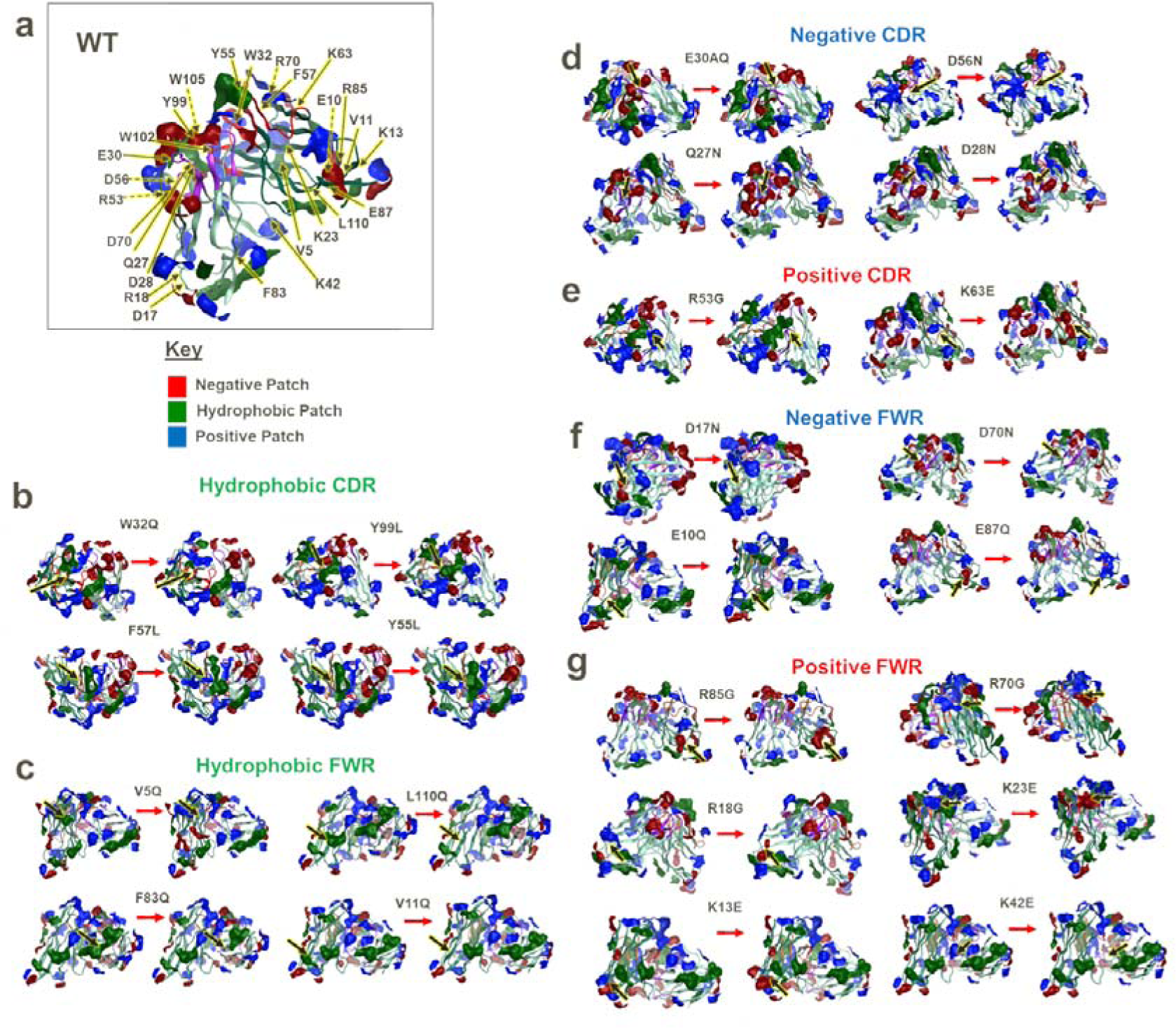
Patch analysis of WT (a) and mutant Fv homology models disrupting hydrophobic patches (green-c-b), negative patches (red-d, f), and positive patches (blue-e, g). VL (light green), VH (dark green), heavy chain CDRs (red), and light chain CDRs (purple) are shown. The WT (left) and corresponding mutant (right) are represented for each molecule. White arrows show the location of the single point mutation. Dashed lines represent residues behind the field of view.

Residues with the highest contributions to positive (blue), negative (red), and hydrophobic (green) patches were identified from patch analysis of the mAb1 WT Fv homology model. Key residues for sequence-based modification included those contributing to positive (blue) (*e.g.,* K42, K23, R18, K13, R85 and R70 for the framework region, and R53 and K63 for CDRs), negative (red) (*e.g.,* D70, E10, E87, D17 for the framework region, and E30A, D56, Q27 and D28 for CDRs) and hydrophobic (green) (*e.g.,* F83, L110, V11, V5 for the framework region, and W32, Y99, F57 and Y55 for CDRs, **Supplementary Tables S1-S2**) patches.

Global patch analysis of mAb1 Fv mutants (**Figure 2**) revealed that R➔G and K➔E mutants (positive patch-targeting*^12^*) exhibited reduced positive patch coverage, while V/W/L/F➔Q and F/Y➔L mutants (hydrophobic patch-targeting*^13^*) showed reduced hydrophobic patch area coverage. D➔N and E➔Q mutants (negative patch-targeting*^14^*) displayed a reduction in negative patch area coverage. However, these mutations did not exclusively impact the targeted patch, with depletion and enhancement of neighbouring patches being observed.

Next, we computed physicochemical molecular descriptors for all candidate mutant Fv homology structures, some of which have shown prior positive or negative correlations with viscosity (**Supplementary Table S3**).^16,17^ We found that charge-based mutant Fvs resulted in changes in predicted net charge, ensemble charge (*ens_charge*), zeta potential, isoelectric point (*pI_seq* and *pI_3D*), and light and heavy chain charge imbalance (*Fv_chml*) (**Supplementary Table S4**). Significant differences in hydrophobicity descriptors were observed with mutants targeting hydrophobic patches (**Supplementary Table S5**).^13^ The therapeutic antibody profiler (TAP)^18,19^ was used to predict developability risk for each candidate mutant (**Supplementary Figure S1**). All mutants were amber-flagged for hydrophobic patches near CDRs, red-flagged for a positive patch targeting mutant (K42E) and a hydrophobic patch targeting mutant (W102Q). We evaluated charge symmetry, with three positive patch-targeting mutants (K42E, R18G and R53G) being amber flagged. From TAP analyses, we identified specific mutants (W102Q, R18G, R53G and K42E) as the ‘*least developable*’ candidates.

### Light chain-heavy chain charge separation

We observed shifts in charge distribution profiles reflected in *Fv_chml* and *FvCSP* descriptors, which indicate charge imbalances between heavy (VH net charge) and light chains (VL net charge). In all cases, VL net charge was negative (-1.23 C for the mAb1 WT) and VH net charge was positive (+3.93 C for the mAb1 WT). Since *FvCSP* is a product of VH and VL charges, we noted a larger difference with negatively-charged VL mutants. For example, with VL and VH charges at -1 C and -4 C, respectively, a 1 C drop in VL net charge reduced *FvCSP* from -4 to -8 C. A 1 C reduction in VH net charge reduced *FvCSP* from -4 C to -3 C. When net charges of either VL or VH chain were 0, *FvCSP* was 0, potentially misinterpreted as no existing charge differences between chains.^20^ This underscores the importance of *Fv_chml* descriptors, which subtract VL charge from VH charge.

Mutants targeting negative patches in VL,^14^ resulted in a ≤0.91 C charge increase, with a similar increase seen for VH mutants. For nearly all VL D➔N mutants, we observed increased *FvCSP* and reduced *Fv_chml*, suggesting enhanced charge symmetry between VH and VL chains. However, VH E➔Q mutants showed a reduction in *FvCSP* and increased *Fv-chml*, indicating increased charge imbalance, absent in Q27N.

Conversely, mutants targeting positive patches^12^ exhibited increased VL negative charge (K42E: -1.9 C), resulting in more negative *FvCSP* and increased *Fv_chml* descriptors. VH mutants had reduced VH charge, approaching VL charge (∼3 C), leading to increased *FvCSP* and decreased *Fv_chml*, reflecting reduced charge imbalance between VL and VH. Mutants primarily targeting hydrophobic patches^13^ resulted in *FvCSP* and *Fv_chml* comparable to mAb1 WT. These data emphasise that single-point mutations in VL versus VH depend on parent WT mAb initial charge symmetry and must be evaluated on a case-by-case basis.

### Triage of candidate mutants

The mAb1 mutant panel was ranked using a summed normalised score, guiding our selection of mutants for expression and physicochemical measurements (**Supplementary Table S7**). We selected two hydrophobic-targeting mutants, four negative patch-targeting mutants, and two positive patch-targeting mutants for expression and subsequent formulation at high concentration (>200 mg/mL). We anticipated that the W32Q mutant, disrupting hydrophobic patches, would significantly reduce viscosity relative to mAb1 WT, while mutants disrupting positive patches (R53G and K42E) would likely show increased viscosity at high concentrations.

### Biophysical Parameters of the Expressed Mutant Panel

We aimed to establish a comprehensive measurement pipeline for the expressed mAb1 mutant panel, correlating these observations with predicted physicochemical descriptors and viscosity-related parameters to understand factors underlying elevated viscosity in high-concentration antibody formulations. We confirmed the sequence identity and post-translational modifications of WT and mutant mAb1 *via* mass spectrometry-based peptide mapping (**Supplementary Table S8**). Additionally, all mutants met the monomeric purity threshold by aSEC (≥ 95%,) (**Supplementary S9**). Apart from W32Q (CDRH2 mutant), mutants retained antigen binding affinity and kinetics equivalent to WT mAb1 (**Supplementary S10**). Next, we analysed the mutants for their hydrophobic, colloidal, electrostatic, and conformational properties.

### Electrostatic properties of the mAb1 mutant panel and the correlation between predicted and experimental parameters

Therapeutic antibodies are typically formulated at high concentrations in the pH 5.2-6.3 range, where the constant regions exhibit a positive net charge, driving repulsive interactions. Variations in charges within the variable region can influence viscosity at high concentrations.^17,21^

Two strategies were employed to generate mutants, targeting positive and negative patches. Therefore, we evaluated electrostatic properties of the mutant mAb1 panel and correlated them with viscosity-concentration profiles. Predicted net charge, isoelectric point (pI), and zeta-potential based on mAb1 Fv (**Supplementary Table S4**) were compared with experimental measurements (**Figure *3***).

**Figure 3.**
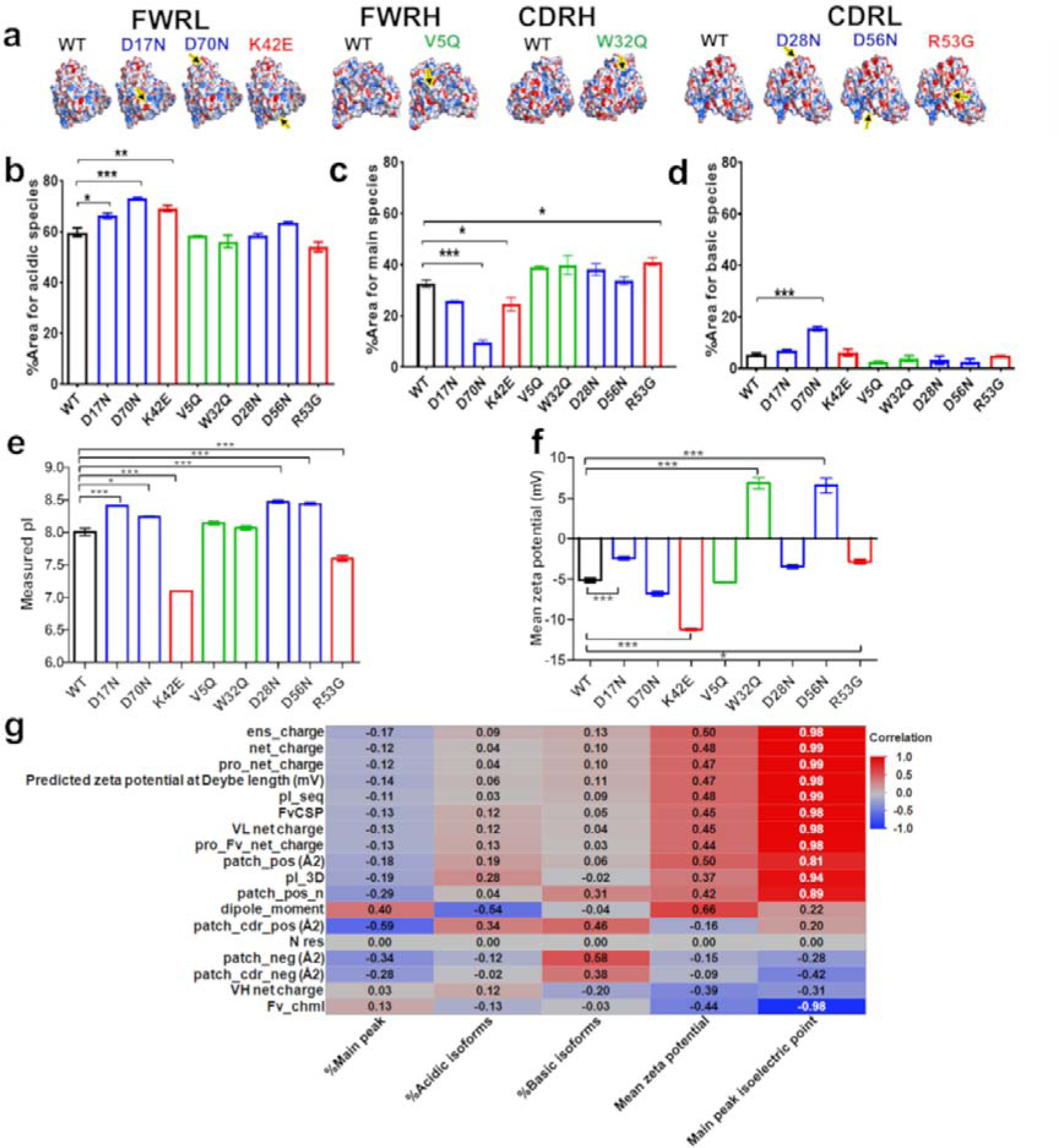
Negative and positive patch disrupting mutants show a strong correlation between predicted and measured PI. **a,** Poisson-Boltzmann surfaces were mapped onto all mAb1 mutant Fv models, categorised by location, demonstrating the impact of single-point mutations on electrostatic distributions around the mutation site (marked by an arrow). **b-f,** Charge-based profiling of mab1 mutant panel with cIEF (N=2) **g,** correlation analyses of zeta-potential showed a weak corelation between the *in silico* descriptor and experimental zeta-potential (N=3) (R=0.47). Strong positive correlations were observed for *pI_seq* and *pI_3D* (sequence and structure based isoelectric point predictions) with the experimental isoelectric points (R=0.99 and 0.94, respectively). A one-way ANOVA with Dunnett’s comparison test was used to compare mAb1 mutants with the WT. *** denotes a P<0.001, ** P<0.01. Non-significant differences are not represented. R values were computed from simple linear regression of *in silico* molecular descriptors and experimental values.

Spatial charge distributions of mutants were visualised with two-dimensional maps (**Supplementary Figure S2** and **Table S6**) to track changes resulting from single point mutations. For example, the D17N mutation led to the loss of a 30 Å^2^ negative patch and a similarly sized hydrophobic patch, with adjacent positive patch surface distributions shifting (WT 2D map numbers 9 and 18 ➔ D17N 2D map numbers 10 and 6). Changes in measured pI were observed, with increased charge for negative patch disrupting mutants, decreased charge for positive patch disrupting mutants, and no significant changes for hydrophobic patch disrupting mutants. (**Figure *3*e**). The majority of mAb1 molecules displayed a negative zeta potential (**Figure *3***f), indicating predominantly attractive forces, except for W32Q and D56N, which had a positive zeta potential. D17N and R53G showed significant increases in zeta potential, while K42E (a positive patch-disrupting mutant) exhibited a reduced zeta potential relative to WT, suggesting increased attractive forces.

We correlated experimental charge data with predicted *in silico* zeta potential and pI descriptors using least-squared polynomial fits (**Figure *3***g). While no correlation was found between the predicted and experimental zeta potential (Pearson correlation coefficient, R =0.47), a strong positive correlation was observed for sequence- and structure-based pI descriptors and measured pI (R=0.99 and 0.94, respectively).

### Hydrophobicity of the Mutant MAb1 Panel and the Correlation Between Predicted and Measured Parameters

As stated above, hydrophobic interactions drive protein-protein interactions and self-association at high formulation concentrations, potentially leading to elevated viscosity. Here, we explored alterations in hydrophobic surface area coverage as a strategy to reduce viscosity, correlating predicted hydrophobicity descriptors with experimental measures.^13^

Using Hydrophobic Interaction Chromatography (HIC), we probed changes in hydrophobicity among the mAb1 mutant panel. We anticipated reduced hydrophobicity for mutants targeting hydrophobic patches, and smaller changes for those targeting charged patches (**Figure 4** and **Supplementary Figure S4**). Indeed, we observed a shorter retention time for W32Q, consistent with predicted reduction in hydrophobicity. Unexpectedly, D70N also showed reduced retention time compared to WT, contrary to predictions. Interestingly, V5Q, predicted to have reduced hydrophobicity, exhibited longer retention time. However, this contradicted predictions, possibly due to differences in targeted hydrophobic patch sizes. Mutants in the CDRL region (D28N, D56N and R53G) showed longer retention times, correlating with spatial hydrophobicity profiles (**Supplementary Figure S2 and Table S6**). Using correlation analysis, we found a strong potential correlation (R=0.87) between normalised hydrophobicity score and summed residue contributions to hydrophobic patch area (*res_hyd*), offering insights into ranking the hydrophobicity of mAb1 mutants.

**Figure 4.**
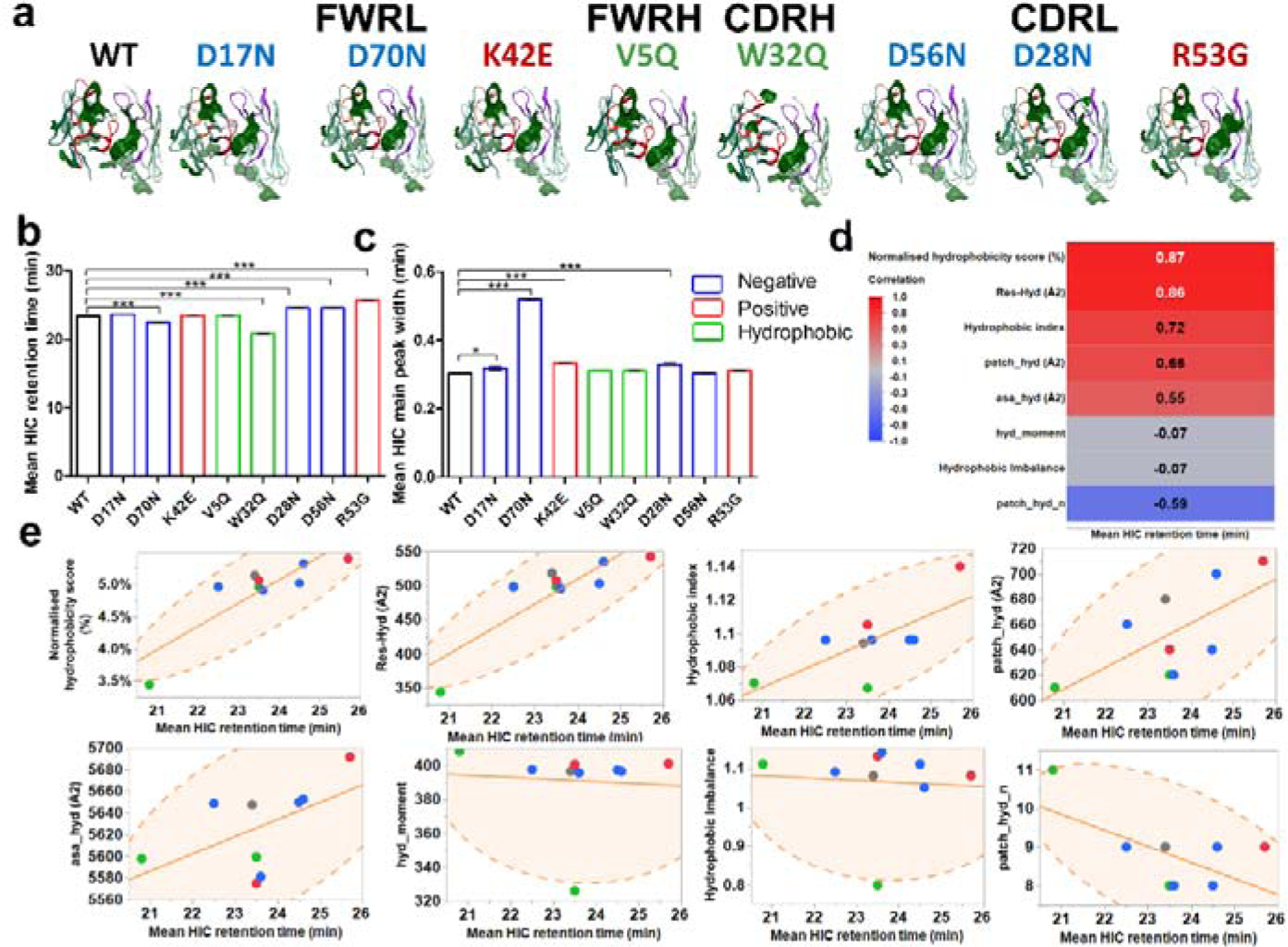
Hydrophobic Interaction Chromatography (HIC) of the WT and mutant mAb1 panel and correlation with predicted hydrophobicity molecular descriptors. **a.** protein patch surface maps are depicted for all mAb1 mutants, filtered for hydrophobic patches (green)**. b,** HIC retention time and **c,** corresponding HIC peak widths for the mAb1 mutants (N=2). Statistical significance was assessed with a one-way ANOVA with Dunnett’s comparison test to WT (*** denotes a P<0.001, * P<0.1). Non-significant differences are not represented. **d,** correlation analysis between *in silico* hydrophobicity descriptors and experimental retention time for mAb1 mutants. Strong correlations (R ≥±0.8) are labelled in white. **e,** scatterplots showing linear correlations for mAb1 mutants with density of ellipses P= 0.95. All antibodies are colour-coded according to mutants targeting positive (**red**), negative (**blue**), and hydrophobic (**green**) patches.

### Conformational Stability of the Mutant Mab1 Panel

We employed intrinsic fluorescence DSF to measure the effects of single-point mutations on mAb1 conformational stability. We used first derivative 350/330 nm ratio traces and scattering traces were used to calculate the unfolding onset temperature (T_onset_), melting temperatures, and the temperature of aggregation onset (T_agg_). Overall, mutants showed comparable thermal stability, except for W32Q and R53G (**Supplementary Figure S5 and Table S11**). W32Q (hydrophobic patch-targeting) exhibited decreased T_onset_, T_agg_ and T_m1,_ suggesting reduced thermal stability. This reduction may stem from the disruption of a large hydrophobic patch (150 Å^2^), critical for stabilising the CDRH2 domain secondary/tertiary structure. R53G (positive patch-disrupting mutant), also showed reduced thermal stability (decreased T_onset_).

### Propensity for interactions promoting self-association

AC-SINS and high throughput diffusion self-interaction parameters (*k_D_*) were used to determine diffusion coefficients (**Supplementary Figure S6**) as surrogate measures of propensity for protein-protein interactions (**Figure *5***).

**Figure 5.**
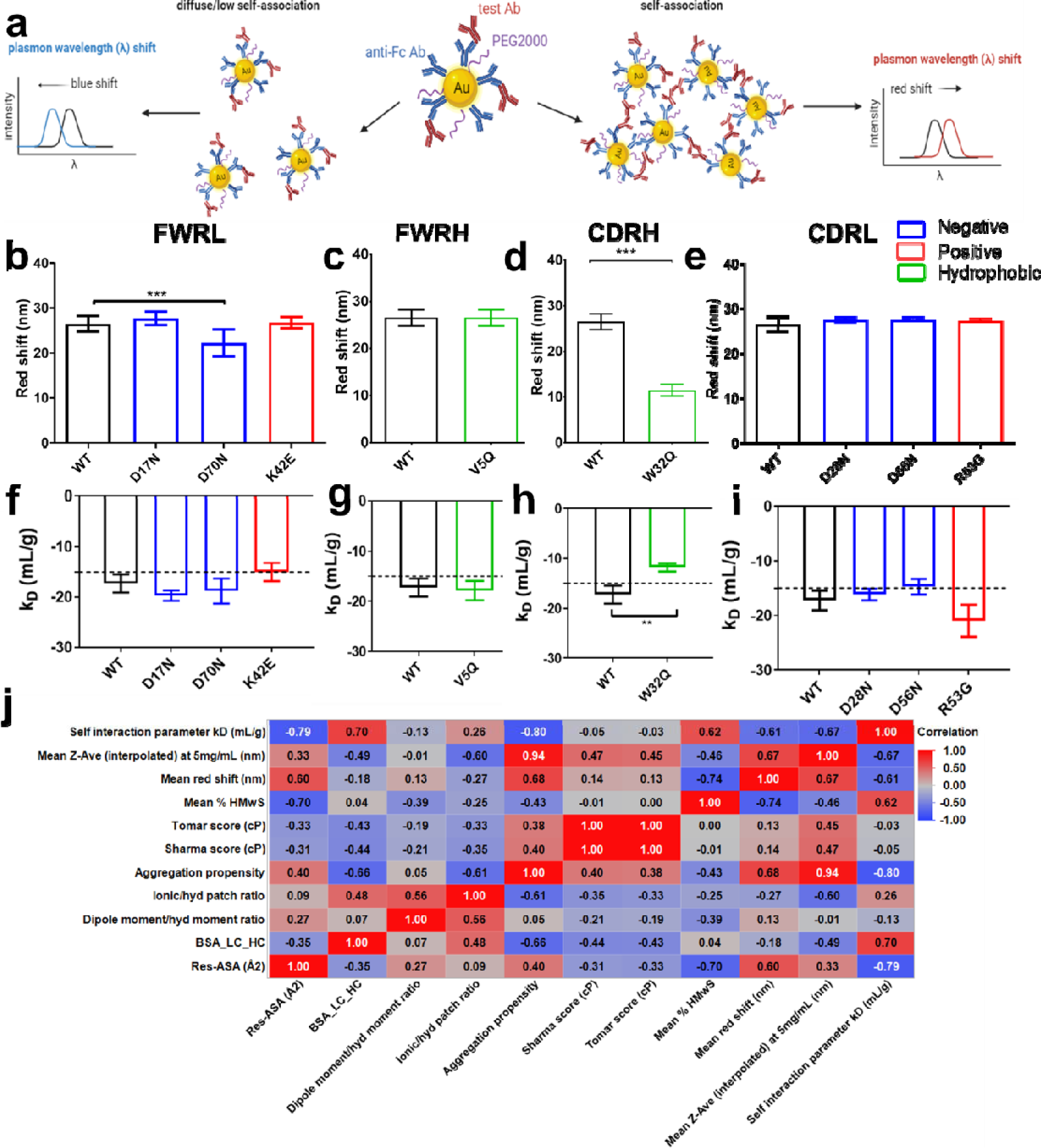
WT and mutant mAb1 panel propensity for self-association as measured with AC-SINS and self-interaction parameter (kD), categorised by mutation location, and coloured by mutation strategy. All antibodies are colour-coded according to mutants targeting positive (**red**), negative (**blue**), and hydrophobic (**green**) patches. **a,** schematic representation of the AC-SINS method and **b-e,** corresponding AC-SINS data (N=4) **f-i,** the self-interaction parameter calculated from analysis of diffusion coefficients (N=2) measured by DLS (1-30 mg/mL). A dotted line at -15 mL/g represents an arbitrary threshold for k_D_. A one-way ANOVA with Dunnett’s comparison test to WT (*** denotes a P<0.001, ** P<0.01). Non-significant differences are not represented. **j,** colloidal interaction experimental results (k_D_ and mean red shift) and hydrodynamic size (Z_ave) and the % high molecular weight species (soluble aggregates) were cross-correlated with *in silico* molecular descriptors describing the structural accessibility (*Res-ASA* and *BSA_LC_HC*), the charge/hydrophobicity ratios, and the aggregation propensity scores. These were selected describe the intrinsic biophysical profile of the mAb1 mutants and their self-interaction propensity. Strong correlations (R ≥±0.8) are labelled in white.

AC-SINS detects self-association by red shifts in UV-Vis spectra (**Figure *5*a**), indicating increased particle size. Compared to the mAb1 WT, D70N and W32Q mutants showed reduced red shift in absorbance measurements (**Figure *5*b, d**), suggesting decreased self-association propensity.

The k_D_ parameter, indicative of protein-protein interaction risk, was comparable to WT for all mutants except W32Q, which had a notably higher k_D_ (>-15 mL/g), signifying reduced short-range attractive self-interactions.^22^ (**Figure *5*h**). D70N showed significant difference in k_D_ compared to WT (**Figure *5*g**). Overall, both AC-SINS and k_D_ data suggest reduced aggregation risk for W32Q.

TANGO aggregation propensity scores, serving as *in* silico predictors of aggregation, negatively correlated with k_D_, soluble aggregates (%HMwS) and hydrodynamic diameter (Z-Ave) (**Figure *5*j**), indicating solvent exposure plays a key role in driving mAb self-association.^23^

### Viscosity-concentration profiles of mAb1 mutants

We evaluated the viscosity of the mAb1 panel at various concentrations using microfluidic rheometry and compared their viscosity profiles to the WT molecule. Non-Newtonian behaviours were not observed across shear sweep experiments (data not shown), so average apparent viscosities were determined with exponential growth fits (**Figure *6*a**). Among the mutants, D70N (negative patch-disrupting FWRL) and W32Q (hydrophobic patch-disrupting CDRH), showed reduced viscosity compared to WT.

**Figure 6.**
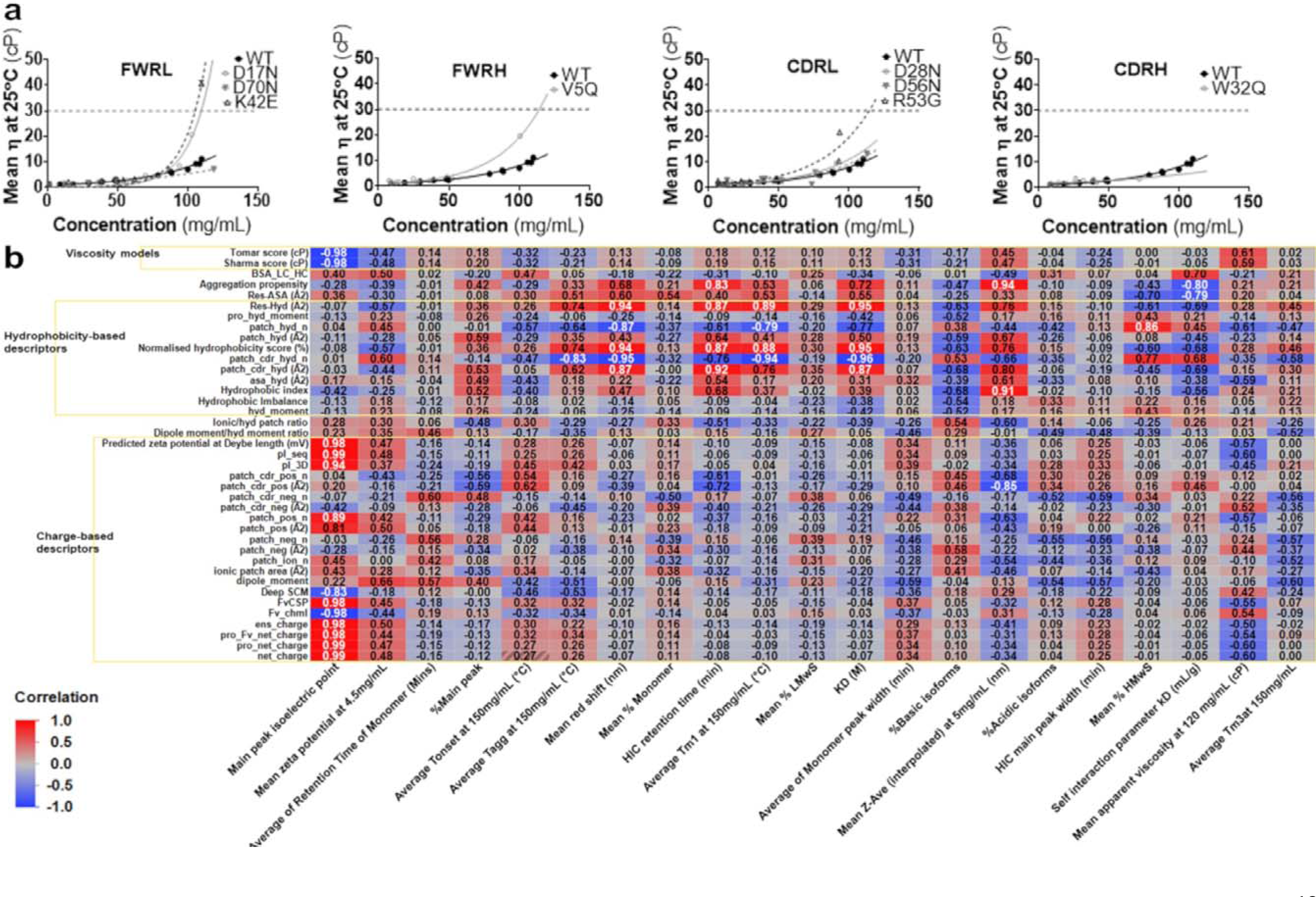
mAb1 mutant panel viscosity correlation heatmap. **a,** Mean apparent viscosity-concentration profiles measured at 25 °C for all mAb1 variants (<120 mg/mL). Dotted grey line at 30cP represents ‘acceptable viscosity’. All measurement data were fitted to exponential growth equations through a least squares fitting method. **b,** correlation heat map values are reported with strong correlations (R >±0.8) in white font.

### Correlating in silico descriptors with biophysical characterisations

We correlated all molecular descriptors used for designing mAb1 mutants with their biophysical characteristics (**Figure *6*b**). For charge-based *in silico* descriptors, the strongest correlations were observed with mean *experimental pI* (**Figure 3**). Weak negative correlations were noted between *net charge* and *pI_seq* and the *mean apparent viscosity* (R= -0.6). A strong negative correlation was found between *patch_cdr_pos area* and the *mean hydrodynamic diameter* (R= -0.85).

Regarding hydrophobicity-based descriptors, strong correlations were observed with *HIC retention time* (**Figure 4**), affinity (KD), AC-SINS red shift and the self-association parameter k_D_. Some strong correlations were also noted between *res_hyd* (R=0.89), *normalised hydrophobicity scores* (R=0.88), and *hydrophobic patch counts* (Fv and near CDRs) (R=-0.94 and -0.79, respectively) with the *Tm_1_ unfolding temperatures*, suggesting the influence of exposed hydrophobic patches on conformational stability of these mAb1 mutants. The number of hydrophobic patches near CDRs was correlated with the *temperature of aggregation onset (T_agg_)*. Additionally, a correlation was observed between the *number of hydrophobic patches* and *the % high molecular weight species* from the aSEC analysis (R=0.86), aligning with hypotheses on the impact of hydrophobic interactions in the mechanism for aggregation.^24,25^ Strong correlations were observed with the TANGO aggregation propensity scores to hydrodynamic diameter (R=0.94), HIC retention time (R=0.83) and k_D_ (R=-0.8). Finally, strong negative correlations were seen with Tomar and Sharma viscosity models, and experimental pIs (-0.98), which is expected as these models are primarily based on charge-related parameters.

## Discussion

The goal of this work was to assess how single-point mutations affect surface exposed electrostatic parameters, hydrophobicity, colloidal, and viscosity behaviour at high formulation concentration in an anti-IL-8 mAb, known for its high solution viscosity at elevated concentrations. We applied three sequence-structure based strategies to design mutants targeting charged (positive and negative) and hydrophobic patches, aiming to compare their effectiveness in predicting and engineering mAb1 developability properties.^26,27^

Our *in silico* predictions of mAb1 physicochemical descriptors revealed notable changes in surface-exposed charged and hydrophobic patches. While mutations in the CDR have previously been associated with reduced mAb viscosity and antigen affinity loss, we expanded our screening to include mutants in the mAb1 heavy and light chain framework regions (**Supplementary Table S10**) ^4,28^ Except for W32Q (a CDRH mutation), all mutants maintained binding affinities for IL-8 equivalent to the WT mAb1 (3.9 nM). W32Q, however, exhibited a five-fold reduction in hydrophobic patch area coverage, suggesting a critical role for tryptophan in antigen binding. This observation aligns with prior studies, where substituting the tryptophan with non-polar and polar amino acids retained binding affinity for phenylalanine mutants, emphasising the importance of the aromatic ring in antigen binding.^29^

We found most mAb1 properties-including monomeric purity and aggregation status-were acceptable for all mAb1 mutants and equivalent to the WT. Overall, point mutations in mAb1 positive and negative patches significantly altered surface potential, inferred colloidal stability, charge heterogeneity and net charge (**Figure *3***).

### Charge-disrupting mutants do not mitigate for elevated viscosity at high-concentration

Adjusting the electrostatic surface potential of mAbs is routinely explored during formulation development, focusing on buffer composition, which alters the excluded volume of the protein in solution (the electroviscous effect).^30,31^ Chow *et al.*^12^ demonstrated viscosity reductions in an IgG4 Fab fragment by reducing charge imbalance across the Fv (R→G and K→E mutants), indicating the impact of positive patch disruption on protein-protein interactions. Conversely, Apgar *et al.*^14^ observed viscosity reduction in mAbs by reducing negative charge, as evidenced by viscosity reduction for D→E to N→Q mutants.^26^

In the current study, the mAb1 WT Fv homology construct exhibited a high proportion of positive patches, indicating a potentially high baseline electrostatic potential with developability risks. We used various *in silico* molecular descriptors (*supplementary information*) to assess developability risks arising from mab1 electrostatic properties. We found negative patch-disrupting mutants reduced charge imbalance^17^, increased net charge,^32^ and ensemble charges,^20^ which have previously been correlated with viscosity reduction. These mutants also exhibited higher pIs, potentially enhancing mAb1 colloidal stability. Conversely, positive patch-disrupting mutants showed reduced ensemble charges and significantly decreased pIs, suggesting diminished colloidal stability.

Zeta potential measurements were conducted on all mAb1 mutants, revealing predominantly negative values consistent with a net negative surface charge observed in the WT mAb1 (pH 6.0). Notably, the K42E mutant exhibited a significantly lower zeta potential compared to the WT, supporting the notion that mutants disrupting positive patches tend to have more negative zeta potentials. Conversely, the W32Q and D56N mutants showed positive zeta potentials, indicating predominant repulsive surface forces associated with hydrophobic and negative patch-disrupting CDR mutants, potentially indicating enhanced colloidal stability. Interestingly, no correlation was found between zeta potential and mutant pI, exemplified by the R53G, which resulted in an increased zeta potential despite having the second-lowest pI value in the mutant panel.

TAP predictions provide charge-based metrics for the mAb1 mutants, with flags indicating charge symmetry primarily in R53G and K42E positive patch targeting mutants (**Supplementary Figure S1**). However, all TAP scores for both positive and negative disrupting mutants fell within an ‘acceptable’ range, suggesting limited discriminatory power of TAP. This lack of differentiation in TAP scores has been noted in previous studies, highlighting potential limitations in its applicability for comprehensive mAb characterization.^2^

### Mutants targeting hydrophobic patches exhibit altered viscosity

Research strategies have explored strategies beyond neutralising charged patches to reduce hydrophobic interactions, for mitigating high concentration mAb stability and viscosity risks.^13^ We computed hydrophobicity-based descriptors for correlation with viscosity and developability, and compared these with HIC retention times (**Figure 4**). Our analyses revealed a reduced hydrophobicity for W32Q, consistent with its predicted decrease in solvent-accessible hydrophobic patch area. However, smaller changes in hydrophobic patch area coverage were undetectable *via* HIC. Mutants with the lowest HIC retention times demonstrated lower solution viscosities (**Figure 6**), indicating a significant role for hydrophobic interactions in driving self-association. Strong correlations were observed between hydrophobic-based *in silico* descriptors and the observed HIC retention times for the mAb1 mutant panel, highlighting the predictive power of these descriptors in understanding viscosity behaviour.

Various research efforts have explored colloidal self-interaction as part of early mAb developability assessments.^33^ The B22 or A2 second virial coefficient and the self-interaction parameter, k_D_, are key metrics capturing the thermodynamic effects of self-associating mAbs at dilute mAb concentrations.^34^ Negative values for B22 and k_D_ indicate attractive protein-protein interactions, associated with decreased formulation stability and increased solution viscosity at high concentrations.^8, 12, 35^ In this study, all mAb1 mutants exhibited negative k_D_ values, with the W32Q mutant showing a less negative k_D_, aligning with its reduced hydrophobicity. The AC-SINS assay further supported reduced self-association propensity for W32Q, consistent with the measured k_D_ (**Figure *5***). Trends were observed between colloidal parameters measured at lower mAb1 concentrations and viscosity-concentration profiles (<120mg/mL), indicating reduced self-association propensities and viscosities for D70N and W32Q.

Most mutants showed similar unfolding temperatures to the WT, except for W32Q, suggesting a critical role for tryptophan in maintaining a large hydrophobic patch in the CDRH2, which impart stability which is lost upon mutation (**Supplementary Figure S5**). This reduced thermal stability also aligns with the observed reduction in antigen binding for W32Q.

Overall trends for each mAb1 molecule in relation to *in silico* physicochemical descriptors and experimental parameters correlated with developability. Kingsbury et al.^2^ correlated multiple *in silico* parameters with opalescence and viscosity for a dataset of 59 commercial mAbs and observed significant clustering with measured pI, effective charge and charge imbalances related to solution behaviour. Overall, we summarise the WT and mAb1 rankings across *in silico* and experimental molecular descriptors in **Figure 7**.

**Figure 7.**
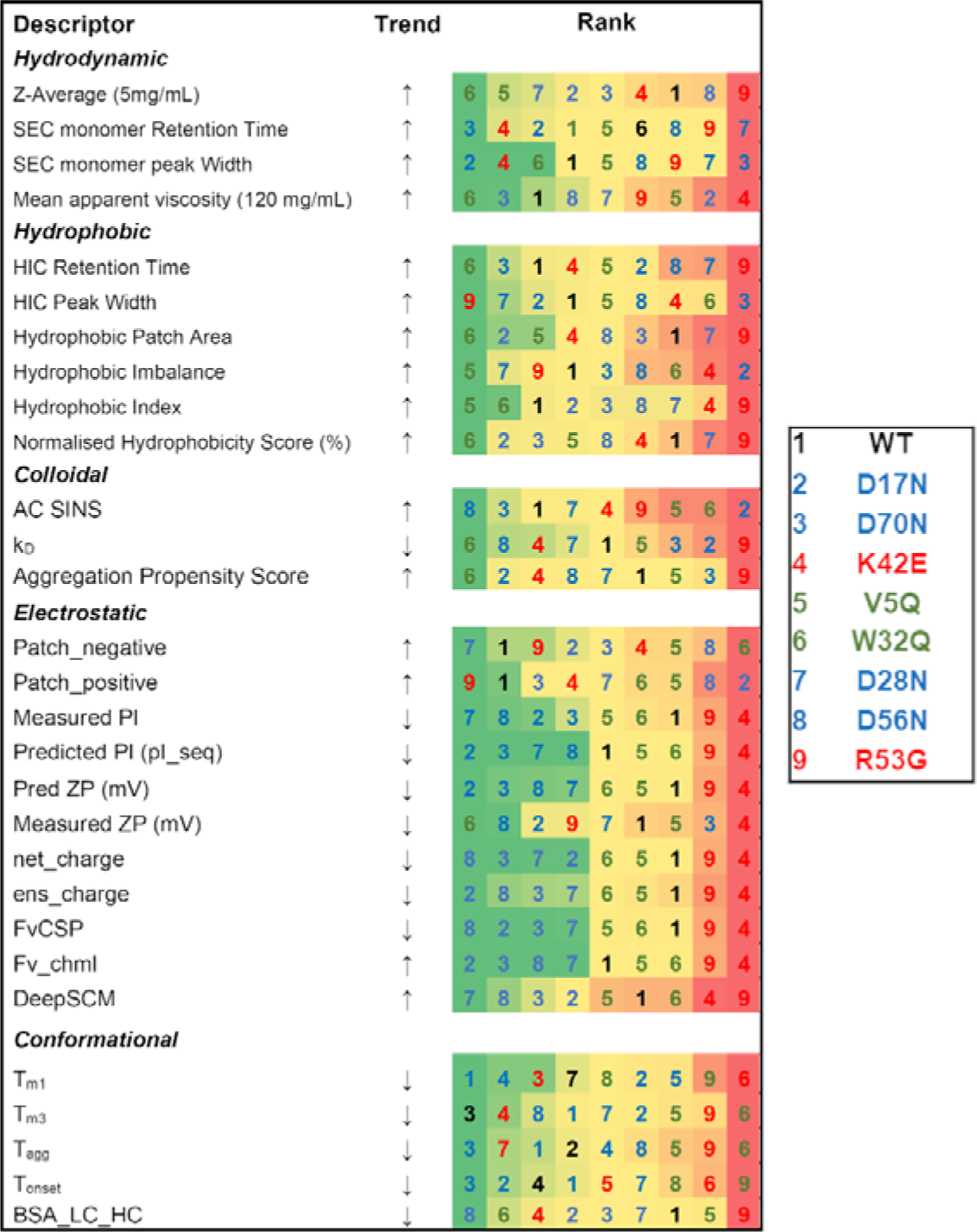
Ranking matrix for the mAb1 mutant panel. A colour-coded from min-max ranking in order of decreasing developability, and categorised by experimental parameters and molecular descriptors.

This is the first study that enables comparison of predictive and empirical approaches to understand the role of electrostatic and hydrophobic patch targeting in altering viscosity in the same mAb molecule. While our findings offer valuable insight into these strategies, there are associated limitations. Unlike previous reports,^9^ we did not observe specific trends in viscosity reduction based on mutation site (CDR versus FWR) in the mAb1 scaffold. Given the variability in charge and non-polar patch distribution among individual mAbs, generalised approaches to reduce molecular interactions driving self-associations may not be suitable and require a systematic design-build-test-learn approach. While we explored single-point mutations, sequence engineering may require multiple mutation sites for improved developability. Previous studies have shown enhanced viscosity reduction through combined substitutions in both VH and VL regions.^36^ Additionally, our computational simulations focused on Fv models and did not consider the influence of hinge and constant domains on biophysical characteristics such as charge and hydrophobicity.

Early-stage assessment of pharmaceutical candidates is crucial for guiding decisions on clinical translation. Various industry-wide criteria are used to triage lead biomolecules, and the use of data-driven sequence-engineering strategies to optimise lead candidates represents a growing field. Our investigation shows that trends observed from molecular descriptors to biophysical properties have a strong dependence on the mutation strategy employed. We find that mutations with significant reductions in hydrophobic patches significantly improved mAb solution viscosity, suggesting the predictive power of hydrophobic-based descriptors. However, mutations altering electrostatic patch coverage alone were insufficient to impact viscosity, irrespective of mutation site. Integrating deep learning approaches holds promise for deeper mechanistic insights into mAb developability, yet challenges such as wider data availability in the pre-competitive research landscape remain. Our study highlights the importance of considering both sequence-based and structural alterations in optimising mAb developability characteristics.

## Materials and Methods

### Computational methods

*In silico* homology modelling and antibody molecular descriptor calculations were performed in the Molecular Operating Environment (MOE) software, version 2020.0901 (Chemical Computing Group, Montreal, Canada).

### Homology modelling

Full sequences of the heavy and light chains of an immunoglobulin G1 (IgG1) wild-type (WT) molecule were inputted as FASTA format into the *MOE sequence editor* and annotated with a Kabat numbering scheme. The *Antibody modeller* in MOE (version 2020.0901) was used to search for similar sequences with solved antibody structures as a template for homology constructs. The variable fragment (Fv) of mAb1 is published as PDB ID: 5OB5 (fAb complex with GroBeta). Fv fragments and full IgG structures were modelled by selecting ‘variable domain’ and ‘immunoglobulin’ model types, respectively. The immunoglobulin model type uses the 1IGY PDB structure as a template to model the Fc region. A refinement gradient limit value of 1 was applied, and C-termini were capped with neutral residues, and superimposed to confirm alignment of structures (**Supplementary Figures S1-S2**). Partial charges were added to all atoms, and energy minimization performed using the AMBER10:EHT default forcefield. The Protein Silo (PSILO) database was used to locate sites of hydrogen bonding and other potential interactions with the GroBeta ligand in complex with the Fv.

### Patch analysis and identification of the mutant panel

The protein patch tool in MOE was applied to the WT Fv homology construct to identify electrostatic and hydrophobic surface patches. To aid visualisation of smaller surface patches, we set the following parameter thresholds: hydrophobic cut-off: ≥0.09 kcal/mol, hydrophobic min area: ≥30 Å^2^, charge cut-off: ≥30 kcal/mol/C, charge min area: ≥30 Å^2^, probe sphere radius: 1.8 Å. The residue contribution to the surface patches was analyzed using the *Protein Properties* tool, selecting the ‘*res_hyd*’, ‘*res_pos*’ and ‘*res_neg*’ descriptors. The top scoring residues were then selected as candidate residues for mutations, excluding terminal residues (**Supplementary Table S1**). Three approaches were implemented to alter solvent-accessible charged patches, by **i)** substituting aromatic hydrophobic residues to leucine or glutamine (L or Q),^13^ and **ii)** substituting positively-charged residues (e.g., N or R) to glutamic acid or glycine (E or G),^12^ and **iii)** substituting negatively-charged glutamic acid or aspartic acid (E or D) to positive residues (e.g., N).^14^ We used *Residue Scan* in MOE to introduce point mutations in the WT mAb1 Fv IgG1 sequence.

### Predicted physicochemical descriptors

We computed a range of physicochemical descriptors (**Supplementary Table S2**) for each Fv model using the MOE *Protein Properties* tool. A NaCl concentration of 0.1 M was used to mimic the ionic strength of the formulation buffer (pH 6). *Hydrophobic imbalance* and *buried surface area*, *Fv_chml* values were generated through BioMOE (version 2021-11-18, Chemical Computing Group, Montreal, Canada) for models protonated to pH 6 using the QuickPrep tool.

### TANGO aggregation propensity

(http://tango.crg.es/tango.jsp).***^37,38^***TANGO aggregation was used to predict the sequence-based propensity for beta-sheet formation for all mutants.

### Ranking mAb1 mutants

Candidate mAb1 mutant variants were ranked using a min-max normalisation method to triage mutants for further investigation. Physicochemical descriptors were selected based on prior correlations with viscosity and weighted evenly. Hydrophobic index, TANGO aggregation propensity, the normalised hydrophobic score (proportion of exposed hydrophobic areas (*Res_hyd*) to the total exposed surface area (*Res_ASA*)), zeta potential, buried surface area between heavy and light chains (*BSA*) and the ensemble charge (*ens_charge*) were parameters used for ranking. Descriptor values were normalised between 0-1 (Equation 3).

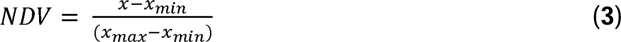

Where *NDV* is the normalised value for a mutant, *x* is the actual descriptor value for a mutant, and *x_min_* and *x_max_* are the minimum and maximum values found in the mutant panel for that descriptor.

A normalised score was calculated by summing each normalised descriptor value (Equation 4.A), or summing 1-normalised descriptor value for descriptors correlating negatively with elevated viscosity (Equation 4.B). Therefore, a lower normalised score overall represented a reduced hypothetical viscosity.

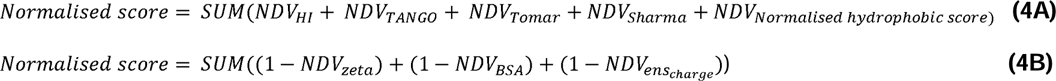

### DeepSCM

(https://github.com/Lailabcode/DeepSCM).^26^ We used the spatial charge map to rank mutants by calculating the charge of side chain atoms of exposed residues of a homology Fv model over molecular dynamics simulations.^26,39^ We inputted mAb1 IgG Fv sequences were inputted as separate heavy and light chain FASTA files and the code was ran in a terminal.

### Protein Expression and Purification

Chinese Hamster Ovary (CHO) K1 GS-KO (glutamine-synthetase-knockout) cells were used for expression of the mAb1 panel. Sequences for mAb1 variants underwent codon optimisation and plasmid generation by Atum Biosciences. The heavy and light chain genes were inserted into a large backbone with a cytomegalovirus (CMV) promoter (Newark, California, USA). Leap-in Transposase® pD2500 vectors with a CMV promoter including glutamine synthetase (for selection) and heavy and light chain insertions were nucleofected into CHO cells. Cells were maintained under selection conditions as stable pooled cultures. A fed-batch production process was employed over 15 days, with glucose and supplementary amino acid feeds added at various intervals. Expression media were harvested and the supernatant clarified by centrifugation at 4 °C (4,000 g for 20 minutes) and sterile-filtered. Protein L chromatography on an ÄKTA Avant 150 system (Cytiva, Danaher, USA) was used for purification, followed by a cation exchange polishing step to achieve ≥95% monomeric purity. The purified mAbs were concentrated, diafiltered and buffer exchanged into formulation buffer containing histidine, trehalose, and arginine (pH 6) to a final concentration of ≥200 mg/mL using the Ambr Crossflow system (Sartorius, Germany).

### Analysis of the WT and mutant mAb1 panel biophysical parameters

### Analysis of mAb1 identity and purity

Peptide mapping was used to confirm the full sequence identity for the mAb1 WT and mutant panel. The monomeric purity of WT and mutant mAb1 variants was analysed by analytical size exclusion chromatography (aSEC) with UV detection (*supplementary information*).

### Hydrophobic interaction chromatography

Hydrophobicity of the mAb1 panel was assessed *via* hydrophobic interaction chromatography (HIC) with UV detection. A PolyLC PolyPROPUL 4.6 x 100 mm column was used on an Agilent 1260 series HPLC (Agilent, California, US). The mobile phase A contained high salt (1.3 M ammonium sulfate) in a potassium phosphate buffer (50 mM, pH 7), with stepwise gradient segments. All samples were analysed at a concentration of 1 mg/mL (with a 5 μL injection) at a flow rate of 0.7 mL/min and detected at 214 and 280 nm wavelengths.

### Zeta potential

A Malvern Zetasizer (Malvern Panalytical, Malvern, UK) with a 633 nm laser was used to measure zeta potential by electrophoretic light scattering. The default settings included an equilibration time of 120s, automatic attenuation and 10-100 measurement runs. A 60-second pause was added between measurements and three technical replicate measurements were performed. Both the WT and mutant mAb1 molecules were prepared in formulation buffer and filtered prior to analysis.

### Diffusion self-interaction parameter

We used a stunner (Unchained Labs, CA, USA) dynamic light scattering setup to measure hydrodynamic size, polydispersity, and the diffusion coefficient for each mAb1 mutant. Data were analysed using the Lunatic & Stunner Client software (version 8.1.0.254). The measurement temperature was set as 25 ℃ and five, 10-second measurements were acquired with a corresponding 1% extinction coefficient of 1.55AU*L/(g*cm) for all samples. Custom dispersant settings were applied (viscosity 1.26 cP and refractive index 1.33 at 20 °C) and all mAbs were prepared in formulation buffer (0.5-20 mg/mL) for WT and mutant variants. The Lunatic & Stunner software (v8.1.0.244) were used for data export, and corresponding diffusion coefficients were used to calculate interaction parameters (k_D_) using linear regression plots.

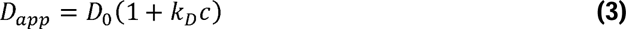

Where D_app_ refers to the apparent diffusion coefficient, D_0_ the self-diffusion coefficient at infinite dilution, and k_D_ the interaction parameter.

### Analysis of mAb1 charge distribution profile

We used the iCE3 capillary isoelectric focusing instrument with a PrinCE autosampler (Protein Simple) to measure charge distribution profiles. A range of pI markers (pI 3.85-8.77) were used to capture all main and impurity isoforms for each sample (Bio-Teche, Protein Simple, USA). To minimise self-association, we used 2M urea and ampholytes (Bio-Teche, Protein Simple, USA) in the pH 3.0-10.0 and 8.0-10.5 ranges at a 1:1 ratio in the buffer mix. All samples were diluted to 1 mg/mL in deionised water prior to a final dilution to 0.4 mg/mL in analyte buffer. The iCE3 instrument was set to the following parameters: a pre-focus voltage of 1,500 V; a 10-12-minute focus voltage of 3,000 V; an autosampler and transfer capillary temperature of 15 °C; UV detection at 280 nm; a sample injection pressure of 2,000 mbar; a pre-focus time of 1 min; and a focus time of 10-12 min. All data were imported to the Empower 3 software (v4, Waters, US) for analysis.

### Analysis of mAb1 self-interaction

We used Affinity-Capture Self-Interaction Nanoparticle Spectroscopy (AC-SINS) to assess self-association propensity in the mAb1 panel. Goat anti-human Fc and whole goat IgG antibodies (Jackson ImmunoResearch, PA, USA) were prepared in 20 mM acetate buffer (pH 4.3) and diluted to achieve final concentrations of 320 µg anti-Fc IgG and 80 µg goat whole IgG, then mixed with 20 nM colloidal gold nanoparticle suspension (Ted Pella Inc., CA, USA, concentration 7.0 x 10^11^ particles /mL). After incubation and centrifugation, mAb1 test samples were prepared at 50 μg/mL in phosphate-buffered saline (Gibco, Thermo Fisher Scientific, MA, USA). Aliquots (99 μL) of each sample were added to wells of a 96-well plate, with 11 μL of gold nanoparticle suspension added to each well, resulting in a final solution concentration of 50 µg/mL test mAb, 10x bead:anti-Fc conjugate and 0.02 mg/mL PEG2000. All samples were mixed, incubated for 90 minutes and gently centrifuged to remove air bubbles. Following incubation, the absorbance spectra (450-650 nm) of the antibody-gold conjugates and analysed using a Pherastar FSX (BMG Labtech Ltd., Germany) plate reader. The spectra were analysed with MARS software (v3.32, BMG Labtech Ltd., Germany), applying smoothing to the best fit curves and the difference in plasmon wavelengths for each sample was calculated. Experimental cutoffs included a <535 nm wavelength for negative controls (*i.e.,* buffer), and a red shift of >10 nm was flagged as a candidate at high risk of self-association.

### Analysis of unfolding temperatures

Thermal differential scanning fluorimetry (DSF) measurements were performed using a Prometheus NT.48 (NanoTemper Technologies, Germany) equipped with back-reflection technology for high-throughput analysis of unfolding temperature (T_m_), calculated from the intrinsic fluorescence intensity ratio of tyrosine and tryptophan (350/330 nm).^40^ Prior to each experiment, the excitation power was set to achieve ≥5,000 counts in the discovery scan. Corresponding profiles were analysed in Prometheus NT.48 and the first derivative calculated. A temperature ramp of 2°C/minute from 20-95 °C was performed for each set of capillaries. Drop lines were assessed and corrected, to determine first-derivative peaks, marking the unfolding temperatures of antibody domains (T_m1_ to T_m3_) and the unfolding onset (T_onset_). The first derivative peak of the scattering profile marked the aggregation temperature (T_agg_) values.

### Viscosity measurement

Viscosity curves were generated using the VROC Initium (Rheosense, United States). The protocol was optimised to measure viscosity samples using the ‘Auto’ shear rate function and fixed shear rates ranging from 100-2000 s^-1^. The resulting data were filtered based on specific criteria, including the exclusion of priming segments, ensuring the percent full scale fell within the 5-95% range, maintaining an R^2^ fit of the pressure sensor position of ≥0.998, and steady plateaus with no drift in transient curves. Exponential-growth decay fits were applied to each viscosity-concentration curve.

### Statistical approaches

GraphPad Prism (v5.04) was used for plotting scatter plots and bar graphs, and ANOVA statistical analysis to determine significant differences in experimental data. JMP Pro (v16.0.0, 2021) was used for the multivariate analyses of computational and experimental data to establish existing correlations.

## Data Availability

The authors declare that all data needed to support the findings of this study are presented in the body of this article and the supplementary information.

## Acknowledgments.

This study was sponsored by GlaxoSmithKline for Georgina Armstrong’s doctoral studies, the UK Engineering and Physical Sciences Research Council (EP/V028960/1), and the UK Biotechnology and Biological Sciences Research Council (BB/Y003268/1). For the purpose of open access, the authors have applied for a CC BY copyright license to any Author Accepted Manuscript version arising from this submission.

