## supplemental for "Enhancing viscosity control in antibody formulations: A framework for the biophysical screening of mutations targeting solvent-accessible hydrophobic and electrostatic patches"

### Supplementary figures and tables

**Patch analysis of candidate mutants**

**Table S1** Top-scoring residues contributing to hydrophobic (*res_hyd*), positive (*res_pos*) and negative (*res_neg*) patches. Residues close to interactions with the GroBeta ligand are marked with an asterisk.

| **res_hyd score (Å2)** | **Residue** | **Position** | **Mutant variant** |
| --- | --- | --- | --- |
| 61.4 | F83 | Framework L | F83Q |
| 57.1 | Y55* | CDRH2 | Y55L |
| 46.9 | L110 | Framework H | L110Q |
| 44.7 | F57* | CDRH2 | F57L |
| 40 | Y99 | CDRH3 | Y99L |
| 33 | V11 | Framework H | V11Q |
| 32.9 | V5 | Framework H | V5Q |
| 28 | W32 | CDRH2 | W32Q |
| **res_pos score (Å2)** | **Residue** | **Position** | **Mutant variant** |
| 52.9 | R53 | CRDL2 | R53G |
| 44.6 | K42 | Framework L | K42E |
| 41.8 | K23 | Framework H | K23E |
| 35.3 | K63 | CDRH2 | K63E |
| 31.8 | R18 | Framework L | R18G |
| 31.3 | K13 | Framework H | K13E |
| 26.3 | R85 | Framework H | R85G |
| 24.8 | R70 | Framework H | R70G |
| **res_neg score (Å2)** | **Residue** | **Position** | **Mutant variant** |
| 70.5 | E30A* | CDRL1 | E30AQ |
| 38.9 | D56 | CDRL2 | D56N |
| 27.1 | Q27 | CDRL1 | Q27N |
| 24.2 | D70 | Framework L | D70N |
| 23.6 | D28 | CDRL1 | D28N |
| 20.3 | E10 | Framework H | E10Q |
| 20.3 | E87 | Framework H | E97Q |
| 18.1 | D17 | Framework L | D17N |

**Patch numbers and corresponding surface area for mAb1 WT and the generated mutants**

**Table S2** Patch numbers and corresponding area coverage for candidate mutant Fv homology constructs.

| Position of mutation | Molecule | patch_ hyd (Å^2^) | patch_ hyd_n | patch_ ion (Å^2^) | patch_ ion_n | patch_ pos (Å^2^) | patch_ pos_n | patch_ neg (Å^2^) | patch_ neg_n | patch_ cdr_pos (Å^2^) | patch_ cdr_pos_n | patch_ cdr_neg (Å^2^) | patch_ cdr_neg_n | patch_ cdr_hyd (Å^2^) | patch_ cdr_hyd_n | Res_ASA (Å^2^) | BSA_ LC_HC |
| --- | --- | --- | --- | --- | --- | --- | --- | --- | --- | --- | --- | --- | --- | --- | --- | --- | --- |
| - | WT | 680 | 9 | 1100 | 23 | 690 | 14 | 410 | 9 | 380 | 6 | 280 | 5 | 280 | 2 | 10078.9 | 681.10 |
| FWR L | D17N | 620 | 8 | 1190 | 23 | 760 | 14 | 430 | 9 | 380 | 6 | 300 | 5 | 260 | 2 | 10109.3 | 681.19 |
| FWR L | D70N | 660 | 9 | 1071 | 22 | 690 | 14 | 380 | 8 | 380 | 6 | 250 | 4 | 260 | 2 | 10051.4 | 681.19 |
| FWR L | F83Q | 540 | 8 | 1190 | 24 | 740 | 14 | 450 | 10 | 430 | 6 | 290 | 5 | 260 | 2 | 10080.5 | 681.19 |
| FWR L | R18G | 660 | 9 | 1080 | 22 | 660 | 13 | 420 | 9 | 380 | 6 | 290 | 5 | 260 | 2 | 10036.4 | 681.19 |
| FWR L | K42E | 640 | 8 | 1060 | 22 | 640 | 13 | 420 | 9 | 380 | 6 | 290 | 5 | 260 | 2 | 10016 | 683.39 |
| FWR H | V5Q | 620 | 8 | 1170 | 24 | 700 | 14 | 470 | 10 | 380 | 6 | 290 | 5 | 260 | 2 | 10060.9 | 680.64 |
| FWR H | E10Q | 660 | 9 | 1140 | 23 | 740 | 15 | 400 | 8 | 380 | 6 | 300 | 5 | 260 | 2 | 10069.2 | 681.19 |
| FWR H | E87Q | 660 | 9 | 1130 | 22 | 740 | 14 | 390 | 8 | 380 | 6 | 290 | 5 | 260 | 2 | 10075.7 | 681.19 |
| FWR H | L110Q | 590 | 8 | 1110 | 23 | 690 | 14 | 420 | 9 | 380 | 6 | 290 | 5 | 260 | 2 | 10058.1 | 681.14 |
| FWR H | V11Q | 620 | 8 | 1120 | 23 | 690 | 14 | 430 | 9 | 380 | 6 | 300 | 5 | 260 | 2 | 10073.1 | 681.19 |
| FWR H | R85G | 660 | 9 | 1070 | 21 | 650 | 13 | 420 | 8 | 380 | 6 | 290 | 5 | 260 | 2 | 10086.4 | 681.19 |
| FWR H | R70G | 630 | 8 | 1080 | 22 | 650 | 13 | 430 | 9 | 340 | 5 | 290 | 5 | 260 | 2 | 10136.5 | 681.19 |
| FWR H | K23E | 620 | 8 | 1140 | 23 | 640 | 13 | 500 | 10 | 380 | 6 | 290 | 5 | 260 | 2 | 10090 | 681.19 |
| FWR H | K13E | 660 | 9 | 1160 | 22 | 660 | 13 | 500 | 9 | 380 | 6 | 290 | 5 | 260 | 2 | 10096.7 | 681.19 |
| CRDL2 | R53G | 710 | 9 | 1030 | 22 | 630 | 13 | 400 | 9 | 320 | 5 | 270 | 5 | 310 | 2 | 10057.6 | 661.34 |
| CDRL2 | D56N | 640 | 8 | 1120 | 22 | 740 | 14 | 380 | 8 | 350 | 5 | 250 | 4 | 270 | 2 | 10032.1 | 702.68 |
| CDRL1 | D28N | 700 | 9 | 1040 | 24 | 690 | 14 | 350 | 10 | 380 | 6 | 220 | 6 | 300 | 2 | 10050.4 | 681.19 |
| CDRL1 | E30aQ | 700 | 9 | 1090 | 24 | 690 | 14 | 400 | 10 | 380 | 6 | 270 | 6 | 300 | 2 | 10038.9 | 684.97 |
| CDRL1 | Q27N | 660 | 9 | 1170 | 22 | 680 | 14 | 490 | 8 | 370 | 6 | 360 | 4 | 260 | 2 | 10055.5 | 681.19 |
| CDRH3 | W105Q | 630 | 8 | 1170 | 24 | 720 | 14 | 450 | 10 | 340 | 5 | 320 | 6 | 260 | 2 | 10096.4 | 682.89 |
| CDRH3 | Y99L | 670 | 9 | 1130 | 23 | 700 | 14 | 430 | 9 | 390 | 6 | 300 | 5 | 270 | 2 | 10072.1 | 661.51 |
| CDRH3 | W102bQ | 670 | 9 | 1110 | 24 | 680 | 14 | 430 | 10 | 370 | 6 | 300 | 6 | 270 | 2 | 10064.5 | 608.28 |
| CDRH2 | W32Q | 610 | 11 | 1120 | 23 | 700 | 14 | 420 | 9 | 390 | 6 | 290 | 5 | 180 | 3 | 9997.4 | 688.06 |
| CDRH2 | F57L | 660 | 9 | 1090 | 23 | 670 | 14 | 420 | 9 | 360 | 6 | 290 | 5 | 260 | 2 | 9999.2 | 697.78 |
| CDRH2 | Y55L | 660 | 9 | 1060 | 22 | 640 | 13 | 420 | 9 | 330 | 5 | 290 | 5 | 260 | 2 | 10036.7 | 678.32 |
| CDRH2 | K63E | 660 | 9 | 1160 | 23 | 640 | 13 | 520 | 10 | 340 | 5 | 390 | 6 | 260 | 2 | 10105.9 | 684.52 |

**Physicochemical molecular descriptors**

**Table S3** Physicochemical descriptors computed for WT and mutant homology models that have been used in previous studies to predict viscosity.

| **Name** | **Description** |
| --- | --- |
| **patch_hyd** | Summed area of hydrophobic patches (Å^2^).^[1]^ |
| **patch_hyd_n** | Summed number of hydrophobic patches.^i^ |
| **patch_pos Å2** | Summed area of positive patches (Å^2^).^i^ |
| **patch_pos_n** | Summed number of positive patches.^i^ |
| **patch_neg Å2** | Summed area of negative patches (Å^2^).^i^ |
| **patch_neg_n** | Summed number of negative patches.^i^ |
| **patch_ion** | Summed area of ionic (positive and negative) patches (Å^2^).^i^ |
| **patch_ion_n** | Summed number of charged (positive and negative) patches.^i^ |
| **asa_hyd Å2** | Solvent-accessible surface area of hydrophobic atoms of a protein (Å^2^).^i^ |
| **patch_cdr_hyd** | Summed area of hydrophobic patches near the CDRs (Å^2^).^i^ |
| **patch_cdr_hyd_n** | Summed number of hydrophobic patches near the CDRs.^i^ |
| **patch_cdr_pos** | Summed area of positive patches near the CDRs (Å^2^).^i^ |
| **patch_cdr_pos_n** | Summed number of positive patches near the CDRs.^i^ |
| **patch_cdr_neg** | Summed area of negative patches near the CDRs (Å^2^).^i^ |
| **patch_cdr_neg_n** | Summed number of negative patches near the CDRs.^i^ |
| **Hydrophobic Imbalance** | A vector that describes the displacement of the superficial geometric centre of the protein when the respective ASA values of each amino acid is considered.^[2]^ This was calculated through the descriptors function in the BioMOE module in MOE 2020. Default parameters were used with no sampling. This was calculated off the Fv model at default values of pH 7.4, temperature of 300K and a salt concentration of 0.1M.^iii^ |
| **Fv_chml** | The Fv heavy chain (V_H_) charge – Fv light chain (V_L_) charge. This was calculated through the descriptors function in the BioMOE module in MOE 2020. Default parameters were used with no sampling. This was calculated from the Fv model at default values of pH 7.4, temperature of 300K and a salt concentration of 0.1M.^[3]^ |
| **Pro_Fv_net_charge** | The protein net charge on Fv only. This was calculated through the descriptors function in the BioMOE module in MOE 2020. Default parameters were used with no sampling. This was calculated off the Fv model at default values of pH 7.4, temperature of 300K and a salt concentration of 0.1M.^iii^ |
| **Pro_net_charge** | The protein net charge. This was calculated through the descriptors function in the BioMOE module in MOE 2020. Default parameters were used with no sampling. This was calculated off the Fv model at default values of pH 7.4, temperature of 300K and a salt concentration of 0.1M.^iii^ |
| **Net_charge** | The formal protein net charge at a given pH. This was calculated through the Protein Properties tool in MOE 2020. The target pH was set to 6, temperature set to 300K and salt concentration to 0.1M.^i^ |
| **Dipole_moment** | Dipole calculated across the protein from uneven distribution of charges. This was calculated through the Protein Properties tool in MOE 2020. The target pH was set to 6, temperature set to 300K and salt concentration to 0.1M.^i^ |
| **Hyd_moment** | Hydrophobicity moment where each residue side chain hydrophobicity is calculated from the Kyte-Doolittle scale across the length of the protein. ^[4]^ This was calculated through the Protein Properties tool in MOE 2020. The target pH was set to 6, temperature set to 300K and salt concentration to 0.1M.^i^ |
| **Hydrophobicity Index** | The summation of hydrophobic residues’ Eisenberg scores over the summation of hydrophilic residues’ Eisenberg scores. Sharma et al. correlated Lower Eisenberg scores with lower viscosity.^[5]^ |
| **Zeta** | Zeta potential is the electrical potential observed at the slipping plane. This was calculated through the Protein Properties tool in MOE 2020. The target pH was set to 6, temperature set to 300K and salt concentration to 0.1M.^i,^ ^[6]^ |
| **pI_seq** | The isoelectric point of a protein calculated from amino acid composition. This was calculated through the Protein Properties tool in MOE 2020. The target pH was set to 6, temperature set to 300K and salt concentration to 0.1M. ^i,^ ^[7]^ |
| **BSA_LC_HC** | The buried surface area (BSA) between the heavy and light chains in Å^2^. This was calculated through the descriptors function in the BioMOE module in MOE 2020. Default parameters were used with no sampling. This was calculated off This was calculated off the Fv model at default values of pH 7.4, temperature of 300K and a salt concentration of 0.1M. ^iii^ |
| **Pro_hyd_moment** | Hydrophobicity moment where each residue side chain hydrophobicity is calculated from the Kyte-Doolittle scale across the length of the protein. This was calculated through the descriptors function in the BioMOE module in MOE 2020. Default parameters were used with no sampling. This was calculated off the Fv model at default values of pH 7.4, temperature of 300K and a salt concentration of 0.1M. ^iii, iv^ |
| **Ens_charge** | The ensemble average charge of the full molecule. This was calculated through the Protein Properties tool in MOE 2020. The target pH was set to 6, temperature set to 300K and salt concentration to 0.1M.^i^ |
| **pI_3D** | The isoelectric point of the molecule calculated through a modified version of Sillero’s model. The PROPKA algorithm is used. This was calculated through the Protein Properties tool in MOE 2020. The target pH was set to 6, temperature set to 300K and salt concentration to 0.1M.^i^ |
| **Fv charge symmetry (FvSCP)** | Charge symmetry of the Fv was calculated with charge of the light chain multiplied by the net charge of the heavy chain.^v^ |
| **Res_ASA** | The summed contribution from each residue to the accessible surface area in Å^2^. This was calculated through the Protein Properties tool in MOE 2020 and manually summed subsequently. The target pH was set to 6, temperature set to 300K and salt concentration to 0.1M.^i^ |
| **Res_hyd** | The summed hydrophobic contribution from each residue to hydrophobic patch area in Å^2^. This was calculated through the Protein Properties tool in MOE 2020 and manually summed subsequently. The target pH was set to 6, temperature set to 300K and salt concentration to 0.1M.^i^ |
| **Dipole moment/hyd moment ratio** | The ratio of Fv dipole moment over the Fv hydrophobic moment to describe the balance of polar versus nonpolar distributions per molecule. This was previously identified as an intrinsic non-redundant descriptor for a dataset of commercial mAbs.^[8]^ |
| **Ionic/hydrophobic patch area ratio** | The ratio of Fv ionic patch area to hydrophobic patch area. This was previously identified as an intrinsic non-redundant descriptor for a dataset of commercial mAbs. ^viii^ |

**Physicochemical descriptor results of mutant variants.**

**Table S4** Charge-based physicochemical descriptors computed for each mAb1 mutant Fv homology construct.

| Position of mutation | Molecule | Fv_chml | pro_Fv_net_charge | pro_net_charge | net_ charge | dipole_moment | Predicted zeta at Deybe length (mV) | pI_seq | ens_ charge | pI_3D | VL net charge | VH net charge | Fv charge symmetry | Deep SCM |
| --- | --- | --- | --- | --- | --- | --- | --- | --- | --- | --- | --- | --- | --- | --- |
| - | WT | 3 | 3.0 | -0.41 | 0.05 | 554.21 | 0.19 | 6.42 | 2.01 | 6.23 | -1.23 | 3.93 | -4.83 | 1197.42 |
| FWR L | D17N | 2 | 4.0 | 0.17 | 0.62 | 578.20 | 1.79 | 6.68 | 3.30 | 7.61 | -0.32 | 3.93 | -1.26 | 1164.43 |
| FWR L | D70N | 2 | 4.0 | 0.17 | 0.63 | 466.75 | 1.58 | 6.68 | 2.90 | 7.61 | -0.32 | 3.93 | -1.26 | 1136.54 |
| FWR L | F83Q | 3 | 3.0 | -0.40 | 0.05 | 554.97 | 0.19 | 6.42 | 2.10 | 6.23 | -1.23 | 3.93 | -4.83 | 1203.71 |
| FWR L | R18G | 4 | 2.0 | -1.29 | -0.84 | 602.97 | -2.18 | 6.07 | 1.31 | 4.92 | -2.18 | 3.91 | -8.52 | 1224.63 |
| FWR L | K42E | 5 | 1.0 | -2.27 | -1.81 | 482.48 | -4.14 | 5.62 | 0.07 | 4.56 | -3.16 | 3.93 | -12.42 | 1253.89 |
| FWR H | V5Q | 3 | 3.0 | -0.41 | 0.05 | 561.51 | 0.19 | 6.42 | 2.25 | 6.23 | -1.23 | 3.83 | -4.71 | 1194.19 |
| FWR H | E10Q | 4 | 4.0 | 0.17 | 0.62 | 614.54 | 1.64 | 6.68 | 3.00 | 7.61 | -1.23 | 4.83 | -5.94 | 1162.09 |
| FWR H | E87Q | 4 | 4.0 | 0.17 | 0.63 | 528.25 | 1.54 | 6.68 | 3.25 | 7.61 | -1.23 | 4.88 | -6.00 | 1178.73 |
| FWR H | L110Q | 3 | 3.0 | -0.41 | 0.05 | 555.58 | 0.19 | 6.42 | 1.96 | 6.23 | -1.23 | 3.91 | -4.81 | 1197.28 |
| FWR H | V11Q | 3 | 3.0 | -0.41 | 0.05 | 557.56 | 0.19 | 6.42 | 2.10 | 6.23 | -1.23 | 3.92 | -4.82 | 1206.02 |
| FWR H | R85G | 2 | 2.0 | -1.30 | -0.84 | 587.77 | -1.98 | 6.07 | 0.98 | 4.92 | -1.23 | 2.93 | -3.60 | 1217.96 |
| FWR H | R70G | 2 | 2.0 | -1.29 | -0.83 | 565.14 | -1.99 | 6.07 | 0.94 | 4.94 | -1.23 | 2.93 | -3.60 | 1261.49 |
| FWR H | K23E | 1 | 1.0 | -2.21 | -1.72 | 390.65 | -4.03 | 5.62 | 0.11 | 4.68 | -1.23 | 0.95 | -1.17 | 1317.79 |
| FWR H | K13E | 1 | 1.0 | -2.21 | -1.73 | 578.86 | -4.40 | 5.62 | 0.25 | 4.64 | -1.23 | 0.94 | -1.16 | 1290.93 |
| CRDL2 | R53G | 4 | 2.0 | -1.29 | -0.83 | 576.62 | -1.96 | 6.07 | 1.01 | 4.93 | -2.18 | 3.93 | -8.57 | 1256.43 |
| CDRL2 | D56N | 2 | 4.0 | 0.17 | 0.63 | 598.31 | 1.51 | 6.68 | 3.00 | 7.61 | -0.32 | 3.82 | -1.22 | 1136.34 |
| CDRL1 | D28N | 2 | 4.0 | 0.16 | 0.62 | 484.86 | 1.46 | 6.68 | 2.88 | 7.58 | -0.34 | 3.93 | -1.34 | 1062.51 |
| CDRL1 | E30aQ | 2 | 4.0 | 0.00 | 0.71 | 511.01 | 1.67 | 6.68 | 2.96 | 7.40 | -0.35 | 3.93 | -1.38 | 1104.96 |
| CDRL1 | Q27N | 3 | 3.0 | -0.47 | 0.05 | 557.86 | 0.20 | 6.42 | 2.11 | 6.23 | -1.23 | 3.93 | -4.83 | 1194.81 |
| CDRH3 | W105Q | 3 | 3.0 | -0.41 | 0.06 | 547.28 | 0.21 | 6.42 | 2.62 | 6.25 | -1.23 | 3.94 | -4.85 | 1188.64 |
| CDRH3 | Y99L | 3 | 3.0 | -0.40 | 0.05 | 551.36 | 0.19 | 6.42 | 1.92 | 6.23 | -1.23 | 3.9 | -4.80 | 1189.43 |
| CDRH3 | W102bQ | 3 | 3.0 | -0.41 | 0.06 | 552.14 | 0.20 | 6.42 | 2.17 | 6.24 | -1.33 | 3.87 | -5.15 | 1190.74 |
| CDRH2 | W32Q | 3 | 3.0 | -0.41 | 0.05 | 566.44 | 0.21 | 6.42 | 2.38 | 6.24 | -1.22 | 3.92 | -4.78 | 1213.87 |
| CDRH2 | F57L | 3 | 3.0 | -0.45 | 0.05 | 555.44 | 0.19 | 6.42 | 2.07 | 6.21 | -1.23 | 3.93 | -4.83 | 1199.18 |
| CDRH2 | Y55L | 3 | 3.0 | -0.41 | 0.05 | 555.09 | 0.19 | 6.42 | 2.11 | 6.23 | -1.23 | 3.93 | -4.83 | 1200.79 |
| CDRH2 | K63E | 1 | 1.0 | -2.19 | -1.71 | 655.50 | -4.03 | 5.62 | 0.13 | 4.65 | -1.23 | 0.94 | -1.16 | 1279.49 |

**Table S5** Hydrophobicity-based physicochemical molecular descriptors and TANGO aggregation propensity scores of mAb1 mutant variants. Dipole and ionic to hydrophobicity ratios are also reported.

| Position of mutation | Molecule | Hydrophobic imbalance | hyd_moment | pro_hyd_moment | Hydrophobic index | Normalised hydrophobicity score (%) | ASA_hyd Å^2^ | Res-Hyd Å^2^ | Dipole moment/hyd patch area | Ionic/hydrophobic patch area ratio | TANGO Aggregation propensity |
| --- | --- | --- | --- | --- | --- | --- | --- | --- | --- | --- | --- |
| - | WT | 1.08 | 396.57 | 396.57 | 1.094 | 5.14 | 5647.53 | 518.50 | 1.07 | 1.62 | 1603.94 |
| FWR L | D17N | 1.14 | 395.70 | 395.70 | 1.096 | 4.91 | 5581.12 | 496 | 1.20 | 1.92 | 1590.12 |
| FWR L | D70N | 1.09 | 397.58 | 397.58 | 1.096 | 4.96 | 5648.77 | 498.3 | 0.95 | 1.62 | 1627.07 |
| FWR L | F83Q | 1.22 | 334.44 | 334.44 | 1.065 | 4.05 | 5575.46 | 408.2 | 1.39 | 2.20 | 1577.65 |
| FWR L | R18G | 1.03 | 432.39 | 432.39 | 1.140 | 4.96 | 5639.26 | 498.1 | 1.23 | 1.64 | 1589.75 |
| FWR L | K42E | 1.13 | 400.56 | 400.56 | 1.105 | 5.06 | 5574.70 | 507.3 | 0.96 | 1.66 | 1602.38 |
| FWR H | V5Q | 0.80 | 325.88 | 325.88 | 1.067 | 4.96 | 5599.16 | 498.8 | 1.15 | 1.89 | 1603.94 |
| FWR H | E10Q | 1.08 | 396.95 | 396.95 | 1.092 | 4.96 | 5662.86 | 499.1 | 1.25 | 1.73 | 1602.71 |
| FWR H | E87Q | 1.09 | 396.28 | 396.28 | 1.092 | 4.94 | 5664.91 | 498 | 1.08 | 1.71 | 1632.24 |
| FWR H | L110Q | 1.03 | 355.72 | 355.72 | 1.067 | 4.22 | 5604.93 | 424.2 | 1.32 | 1.88 | 1530.25 |
| FWR H | V11Q | 1.08 | 378.72 | 378.72 | 1.067 | 4.95 | 5598.55 | 498.6 | 1.14 | 1.81 | 1603.94 |
| FWR H | R85G | 1.07 | 394.59 | 394.59 | 1.140 | 4.94 | 5647.25 | 498.4 | 1.20 | 1.62 | 1603.62 |
| FWR H | R70G | 1.12 | 401.40 | 401.40 | 1.140 | 4.92 | 5666.67 | 498.8 | 1.15 | 1.71 | 1603.94 |
| FWR H | K23E | 1.08 | 399.60 | 399.60 | 1.105 | 4.95 | 5557.77 | 499.4 | 0.80 | 1.84 | 1624.98 |
| FWR H | K13E | 1.08 | 396.35 | 396.35 | 1.105 | 4.94 | 5627.99 | 499.1 | 1.16 | 1.76 | 1605.15 |
| CRDL2 | R53G | 1.08 | 401.05 | 401.05 | 1.140 | 5.40 | 5691.33 | 542.8 | 1.07 | 1.45 | 1897.13 |
| CDRL2 | D56N | 1.11 | 397.34 | 397.34 | 1.096 | 5.02 | 5649.88 | 503.3 | 1.20 | 1.75 | 1602.95 |
| CDRL1 | D28N | 1.05 | 396.57 | 396.57 | 1.096 | 5.32 | 5652.63 | 535 | 0.91 | 1.49 | 1603.43 |
| CDRL1 | E30aQ | 1.06 | 397.01 | 397.01 | 1.092 | 5.37 | 5655.14 | 538.6 | 0.96 | 1.56 | 1638 |
| CDRL1 | Q27N | 1.10 | 396.47 | 396.47 | 1.095 | 4.98 | 5618.84 | 500.3 | 1.12 | 1.77 | 1603.79 |
| CDRH3 | W105Q | 1.03 | 380.86 | 380.86 | 1.070 | 4.94 | 5599.76 | 498.6 | 1.09 | 1.86 | 1603.88 |
| CDRH3 | Y99L | 1.09 | 393.52 | 393.52 | 1.105 | 5.11 | 5675.74 | 514.2 | 1.08 | 1.69 | 1603.96 |
| CDRH3 | W102bQ | 1.04 | 402.35 | 402.35 | 1.070 | 4.97 | 5598.06 | 500.3 | 1.10 | 1.66 | 1604.05 |
| CDRH2 | W32Q | 1.11 | 408.27 | 408.27 | 1.070 | 3.44 | 5597.72 | 343.7 | 1.67 | 1.84 | 1357.71 |
| CDRH2 | F57L | 1.13 | 387.53 | 387.53 | 1.092 | 4.91 | 5663.06 | 491.3 | 1.13 | 1.65 | 1602.02 |
| CDRH2 | Y55L | 1.11 | 353.42 | 353.42 | 1.105 | 4.98 | 5664.70 | 500.3 | 1.13 | 1.61 | 1605.53 |
| CDRH2 | K63E | 1.08 | 392.53 | 392.53 | 1.105 | 4.93 | 5608.40 | 498.3 | 1.34 | 1.76 | 1605.17 |

**Therapeutic antibody profiler (TAP) scores for mAb1 mutant variants**

***Therapeutic Antibody Profiler*** (<https://opig.stats.ox.ac.uk/webapps/sabdab-sabpred/sabpred/tap>). The therapeutic antibody profiler (TAP) is a developability ranking tool that incorporates CDR length, hydrophobicity, positive and negative charges of CDR patches and Fv charge symmetry of homology structures.***^18^*** The web application was used to submit heavy and light chain sequences of the wild-type and mutant panel.

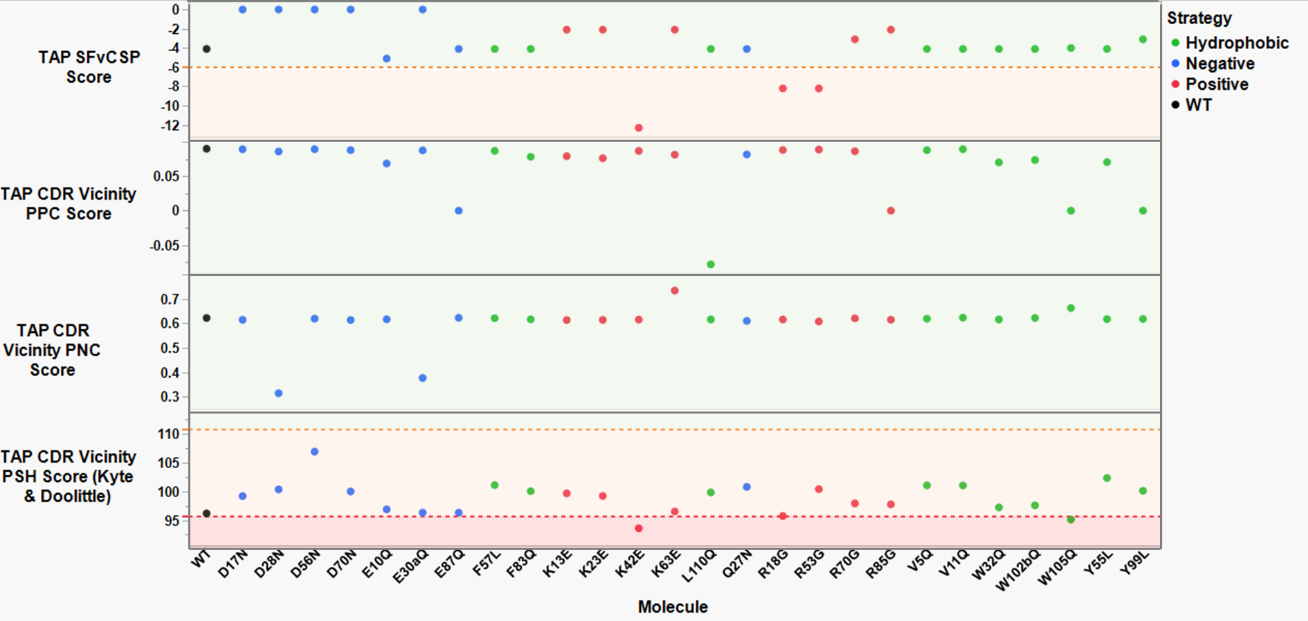

**Figure S1** The Therapeutic Antibody Profiler (TAP) tool computed four structural attributes for mAb1 candidate mutants. This tool used the heavy and light chain sequences of the variable regions for each molecule and the ABodyBuilder2 tool was used to construct homology models. The red-amber-green thresholds were set from previous work analysing 137 clinical stage antibodies. For all mutants the CDR length was 46 residues which was within the green threshold. Structural Fv Charge Symmetry Parameter (SFvCSP) showed three positive-patch disrupting mutants with amber flags (K42E, R18G and R53G). The patches of positive charge (PPC) metric and the patches of negative charge (PNC) metric across the CDR vicinity showed no flags for all mutants. However, all mutants had at least an amber flag for the patches of Surface Hydrophobicity (PSH) metric across the CDR vicinity, with two red flags for K42E and W105Q.

Modified Fv patch areas of expressed mutants

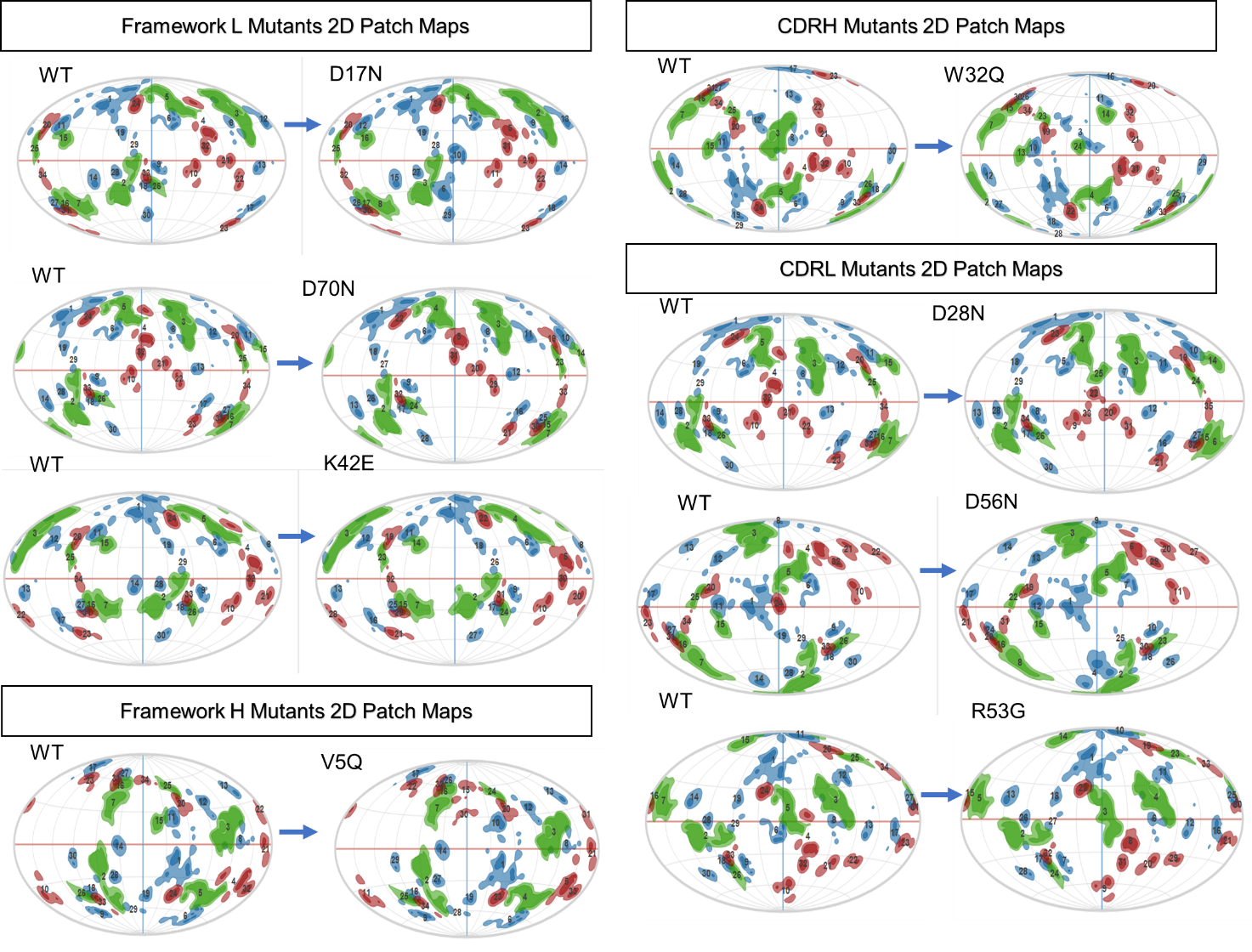

**Figure S2** *Two-dimensional patch maps of expressed mutants Fv homology constructs*. Hydrophobic (green), positive (blue) and negative (red) patches were analysed for the area and energy changes for each expressed mAb1 mutant. The field of view is rotated for each WT and expressed mutant pair to have the site of mutation at the centre.

**Table S6** Quantitation of specific modified patch areas and energy changes for expressed mutants Fv homology constructs.

| **Molecule** | **WT 2D Map Number** | **Patch Type** | **Patch Area**  (Å^2^) | | **Average energy per Å^2^**  (kcal/mol) | | **Other significant residues** |
| --- | --- | --- | --- | --- | --- | --- | --- |
|  |  |  | **WT** | **Mutant** | **WT** | **Mutant** |  |
| **D17N** | 33 | neg | 30 | Removed | -49.42 | Removed | S14, G16 |
|  | 26 | hyd | 30 | Removed | -0.11 | Removed | P8, L11, A13, V19 |
| **D70N** | 10 | neg | 60 | Removed | -49.72 | Removed | Q24, Q69 |
| **K42E** | 14 | pos | 50 | Removed | -54.26 | Removed | P40, G41 |
|  | 28 | pos | 30 | Removed | -51.72 | Removed | K39, P40, E81, F83 |
| **W32Q** | 3 | hyd | 150 | 30 | -0.16 | -0.13 | Y51C |
|  | 8 | pos | 70 | 130 | -41.71 | -48.24 | H91, E93, S93A, P95 |
| **V5Q** | 15 | hyd | 40 | Removed | -0.14 | Removed | K23 |
|  | 11 | pos | 50 | 60 | -61.18 | -61.1 | K23 T74 S75 |
| **D28N** | 32 | neg | 30 | 30 | -55.96 | -43.85 | E30, G68 |
|  | N/A | hyd | 30 | Addition | -0.15 | Addition | E30A, Y32 |
| **D56N** | 24 | neg | 40 | Removed | -69.48 | Removed | N/A |
|  | 19 | pos | 40 | Removed | -45.54 | Removed | K45, G57, V58, P59 |
| **R53G** | 6 | pos | 80 | Removed | -53.93 | Removed | Y50 T52 L54 |
|  | 5 | hyd | 110 | 150 | -0.15 | -0.16 | Y32, Y49, Y50 |

**Triage of candidate mutants**

**Table S7** Top and bottom scoring mAb1 mutants progressed to experimental characterisation based on min-max normalisation. Scoring was based on hydrophobic index, zeta potential, BSA_LC_HC, ens_charge, normalised hydrophobicity, and TANGO aggregation propensity. Each descriptor value was weighted evenly and normalised to ensure that the lower the score, the increased likelihood of reduced hypothesised viscosity.

| **Molecule** | **Mutation** | **Summed normalised score** |
| --- | --- | --- |
| WT (-) | - | 3.28 |
| W32Q (CDRH2) | Hydrophobic | 1.76 |
| D56N (CDRL2) | Negative | 2.13 |
| D17N (FWL) | Negative | 2.21 |
| D70N (FWL) | Negative | 2.35 |
| V5Q (FWH) | Hydrophobic | 2.37 |
| D28N (CDRL1) | Negative | 2.42 |
| R53G (CDRL2) | Positive | 5.83 |
| K42E (FWL) | Positive | 6.22 |

**Biophysical Characterisation**

*Analysis of identity by mass spectrometry*

The sequence and composition of the mAb1 panel was verified using peptide fingerprinting mass spectrometry. 250μg of each sample was denatured with guanidine buffer (6M, pH 7.5), reduced with dithiothreitol (DTT 1M) and incubated for 20 minutes at ambient temperature. All samples were alkylated with 1M sodium iodoacetate and incubated for a further 30 minutes at ambient temperature and protected from light. A further reduction step was performed in DTT (1M), and the samples were desalted using Micro Bio-Spin 6 size exclusion columns (Bio-Rad, CA, USA). Samples were enzyme-digested with either trypsin or chymotrypsin (both sequencing-grade, Promega, WI, USA) at a 1:20 (w/w) ratio of chymotrypsin: mAb in a digestion buffer containing 50mM Tris, 1mM calcium chloride dihydrate (pH 7.5). Samples were incubated at 37 °C under agitation for two hours, prior to liquid chromatography-mass spectrometry (LC-MS) analysis with an Orbitrap Exploris™ 240 Mass Spectrometer (Thermo Fisher Scientific, MA, USA), controlled by Xcalibur software (version 4.4.16.14, Thermo Fisher Scientific, MA, USA). An ACQUITY UPLC PEPTIDE CSH C18 (Waters, US) 1.7 µm, 2.1 mm x 150 mm column was used for separating digested peptides with a column temperature of 40 °C. Mobile phase A was 0.1% Formic Acid LC-MS grade (Thermo Fisher Scientific, MA, USA) in LC-MS grade water and B was 0.1% Formic Acid in Acetonitrile LC-MS grade (Thermo Fisher Scientific, MA, USA). Step wise gradients were applied with 5-40 %B (over 80 min), 40-100 %B (5 min), plateau of 100 %B (5 min), and a return to 5 %B (10 min). The flow rate was set at 200 μL/min and the UV was monitored at 214 nm.

The Orbitrap Exploris 240 MS system was operated in the positive ion mode. Tandem MS/MS analyses were performed for the identification of peptide in data dependent mode. Full MS scan data acquired within a 200-2000 m/z scan range, 60,000 resolution over 100ms injection time, followed by 5 sequential MS/MS scan with orbitrap resolution target of 15000. A minimum intensity threshold was set to 1000 with a custom dynamic exclusion filter applied. Charge states were filtered to include charges of 2-8 and the number of dependent scans was set to 5. A 2 m/z isolation window was applied for the ddMS scan with HCD collision energies set to 20, 25 and 30% over 200 ms injection time. MS2 data acquired in profile mode.The MS2 AGC target was set at 100% whereas full scan AGC target was set at 300%. Byos software (version 5.0-88 (2022.12), Protein Metrics, CA, USA) was used to processing of peptide fragments using the following parameters: Precursor Mass Tolerance set at 20 ppm, Fragment Mass Tolerance 1 and 2 set at 20 ppm, Cleavage Site(s) set as RK (trypsin) and WFLY (chymotrypsin), Missed Cleavages set at 2, Cleavage Side set as C-terminal and Fragmentation type set as QTOF/HCD. The post translation modifications (PTMs) screened for were methylation, oxidation, deamidation and pyroglutamate formation.

**Table S8** Verification of mAb1 WT and mutant variant identity by peptide fragmentations. Trypsin or chymotrypsin digest of this peptide following the same methodology showed coverage of this missing peptide, ensuring full identity verification. For post-translational modifications (PTMs), the % detection was relative to only peptides with expected full enzyme cleavage. PTMs with relative detection were noted. *HC: Heavy chain; LC: Light chain; mwt: molecular weight; PTM: post-translational modification*

| **Molecule** | **LC coverage (%)** | **HC coverage (%)** | **LC mwt (Da)** | **HC mwt (Da)** | **LC PTMs** | **HC PTMs** |
| --- | --- | --- | --- | --- | --- | --- |
| WT | 97.66 | 96.66 | 23433.83 | 49204.09 | M4 oxidation (0.4%) | M81 oxidation (0.2%),  N317 deamidation (0.6%), M254 oxidation (4.2%), N363 deamidation (0.4%), M430 oxidation (1.8%) |
| D17N | 97.66 | 97.44 | 23432.85 | 49204.09 | M4 oxidation (0.3%) | M81 oxidation (0.2%), M254 oxidation (4.5%), N317 deamidation (0.1%), N363 deamidation (0.4%), M430 oxidation (2.1%) |
| D70N | 99.07 | 98.22 | 23432.85 | 49204.09 | M4 oxidation (0.3%), p*ossible N70 deamidation but not confirmed due to poor fragmentation (see map coverage below Figure S3)* | M81 oxidation (0.2%), M254 oxidation (4.8%), N288 deamidation (0.1%), N317 deamidation (0.3%), N363 deamidation (0.2%), N436 deamidation (1.42%) |
| K42E | 97.66 | 98.22 | 23434.77 | 49204.09 | M4 oxidation (0.3%) | M81 oxidation (1.12%), N317 deamidation (0.4%), M430 oxidation (1.9%), N436 deamidation (3.3%) |
| V5Q | 97.66 | 98.22 | 23433.83 | 49233.09 | None | N363 deamidation (0.2%) |
| W32Q | 97.66 | 79.73* | 23433.83 | 49146.01 | None | S methylation (100%), N317 deamidation (2.5%) |
| D28N | 97.66 | 98.22 | 23432.85 | 49204.09 | N28 deamidation (12.3%) | N317 deamidation (0.4%), N363 deamidation (0.2%), N436 deamidation (1%) |
| D56N | 97.66 | 98.22 | 23432.85 | 49204.09 | N56 deamidation (1.85%) | N436 deamidation (0.8%) |
| R53G | 97.66 | 98.22 | 23334.7 | 49204.09 | M4 oxidation (1%) | M430 oxidation (1.6%) |

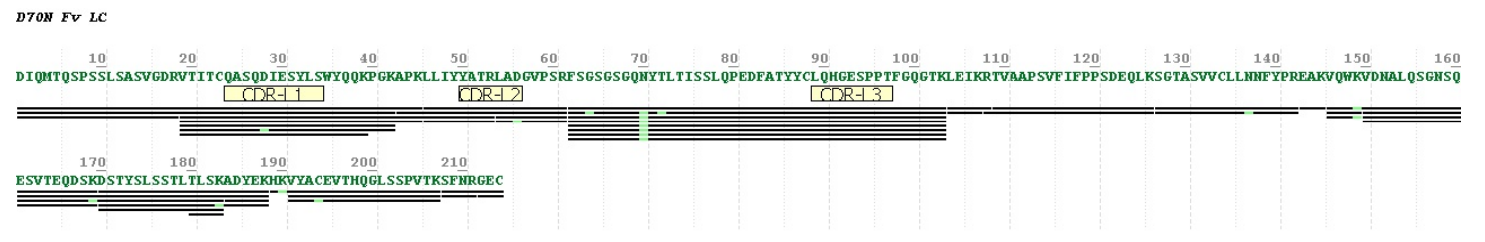

**Figure S3** Peptide map coverage for D70N light chain. Possible modification site at N70 flagged (light green).

***Analysis of Monomeric Purity by Analytical Size Exclusion Chromatography (aSEC)***

Samples were injected onto a TSKgel Super SW3000, 4.6 x 300 mm (TOSOH Bioscience, United States) column on an Agilent 1260 series HPLC, with 0.1M sodium phosphate containing 400 mM NaCl (pH 6.8) as the mobile phase. All samples were analysed at 5 mg/mL at a 0.2 mL/min flow rate, and detected at 280 nm. The OpenLab CDS Data Analysis software (version 2.6, Agilent, California, US) was used to process and integrate the chromatograms. Areas under the chromatographic peaks were integrated to quantify the monomeric mAb, and high and low molecular weight species. The target monomeric purity of ≥95% was met by all WT and mutant mAb1 molecules and aSEC was used to monitor physicochemical stability, by monitoring changes in chromatogram peak retention times and profiles for each molecule. Analysis of the expressed mAb1 mutants showed retention times comparable to the mAb1 WT IgG1 (~27.5 minutes), except for the D70N mutant, which had a consistent reduced retention time of ~26.6 minutes suggesting a slight increase in molecular size.

**Table S9** Monomeric purity of all mAb1 molecules (N=3).

| Mab | RT (min) | Peak Width (min) | %HMW species | %Monomer | % LMW species |
| --- | --- | --- | --- | --- | --- |
| *WT* | 27.5 (±0.5) | 0.53 (±0.05) | 1.3 (±0.2) | 97.1 (±0.5) | 1.6 (±0.5) |
| *D17N (FWL)* | 27.4 (±0.3) | 0.51 (±0.05) | 1.2 (±0.1) | 98.0 (±0.5) | 0.9 (±0.5) |
| *D70N (FWL)* | 26.6 (±0.3) | 0.75 (±0.03) | 1.5 (±0.2) | 98.0 (±0.4) | 0.6 (±0.3) |
| *K42E (FWL)* | 27.3 (±0.3) | 0.51 (±0.05) | 1.4 (±0.2) | 97.2 (±0.6) | 1.4 (±0.5) |
| *V5Q (FWH)* | 27.5 (±0.5) | 0.53 (±0.06) | 1.2 (±0.3) | 97.4 (±0.7) | 1.4 (±0.5) |
| *W32Q (CDRH2)* | 27.5 (±0.6) | 0.51 (±0.05) | 2.1 (±0.6) | 96.8 (±1.1) | 1.1 (±0.5) |
| *D28N (CDRL1)* | 27.6 (±0.5) | 0.55 (±0.1) | 1.8 (±0.3) | 96.5 (±0.8) | 1.7 (±0.8) |
| *D56N (CDRL2)* | 27.5 (±0.5) | 0.53 (±0.05) | 1.4 (±0.6) | 97.0 (±1.1) | 1.7 (±0.5) |
| *R53G (CDRL1)* | 27.5 (±0.5) | 0.53 (±0.05) | 1.5 (±0.4) | 97.2 (±0.7) | 1.2 (±0.4) |

***Hydrophobic interaction chromatography***

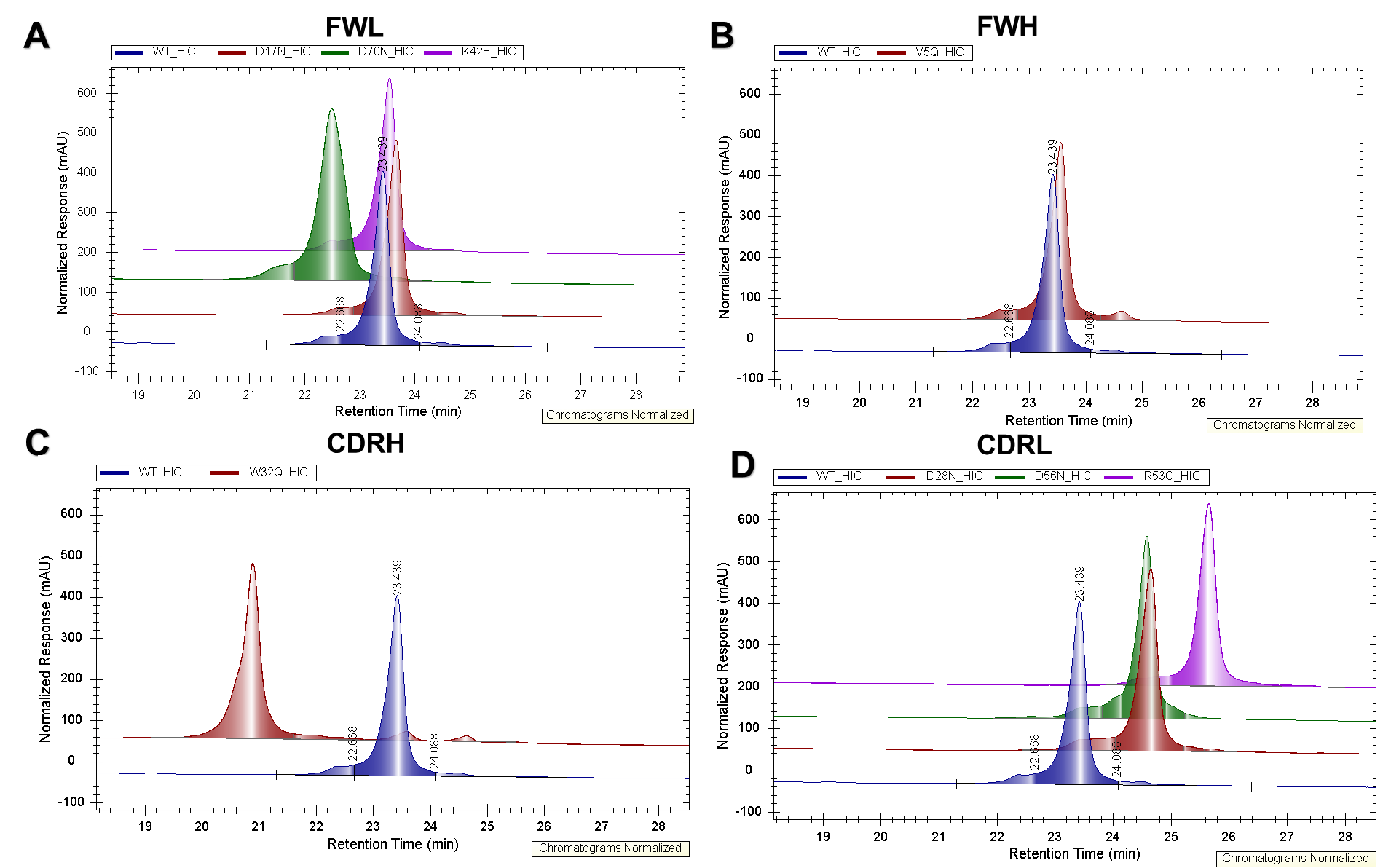

**Figure S4** Hydrophobic interaction chromatography chromatograms for mAb1 mutants. For all chromatograms, the WT (blue) retention time was observed at ~23.5 minutes. For FWL mutants (A), only D70N (green) had a shift in retention time and an increase in peak breadth. A slight increase in retention time was seen for FWH (B) V5Q mutant (red), and a large reduction in retention time for the CDRH (C) W32Q mutant (red), demonstrating the impact of size of the original hydrophobic patch targeted. For all CDRL mutants (D), an increased retention time was observed relative to WT, particularly for R53G (pink) where there was a predicted increase in neighbouring hydrophobic patch size from this disruption of a positive patch.

**Binding analysis**

We used a Biacore 8K+ surface plasmon resonance (Cytiva, Danaher, USA) to compare the association and dissociation rates, and affinity for to IL-8 carrier-free antigen (R&D systems, USA) between the WT and mutant panel. IL-8 (0.5 µg/mL) was immobilised onto one flow cell of a Biacore CM3 dextran chip (Cytiva, Danaher, USA). The experiment consisted of ten start-up cycles, followed by ten antibody injections at a flow rate of 30 μL/min and a temperature of 25 °C. The contact time was 240 seconds, and dissociation was monitored over 900 seconds after injection. All antibodies (0.31-20 µg/mL) were formulated in phosphate-buffered saline containing 0.05% Tween 20, with the same running buffer composition. Surfaces were regenerated between measurements using 10 mM glycine (pH 1.5) and 3 M guanidine. The data were analyzed using Biacore Insight Evaluation software (version 4.0.8.20368, Cytiva, USA) with a 1:1 Langmuir binding model.

To determine the apparent dissociation (ka) and dissociation rate constants (kd). The $\frac{k_{d}}{k_{a}}$ ratio was used to determine the equilibrium dissociation constant (KD).

The impact of introducing single-point mutations on the ligand binding affinity of mAb1 mutant variants was measured by SPR. The mean binding affinity across all mutant variants was equivalent to the mAb1 WT (3.92 nM), except for the W32Q mutant (CDRH), which had no binding affinity for the target antigen.

**Table S10** Biacore analysis of binding kinetics. Wild-type and mutant mAb1 binding to an IL-8 antigen was assessed with SPR. Data in the table (all from 2 replicates) includes the binding on-rate (k_a_), the binding off-rate (k_d_) and the equilibrium dissociation constant (KD), as well as the maximum response (R_max_) and goodness of fit (Chi-squared) of the 1:1 binding model. All framework mutants and CDRL mutants showed no significant change in affinity relative to the mAb1 WT. The WàQ single point mutation in the CDRH2 domain knocked out all binding affinity to IL8 antigen. FWL: light chain framework region; FWH: heavy chain framework region; CDRH2: heavy chain complementarity-determining region 2; CDRL1: light chain complementarity-determining region 1; CDRL2: light chain complementarity-determining region 2 (N=2).

| Molecule | 1:1 binding kinetics | | | | Kinetics (Χ^2^) |
| --- | --- | --- | --- | --- | --- |
|  | **k_a_ x10^5^**  (M^-1^s^-1^) | **k_d_ x10^-4^** (s^-1^) | **KD** (nM) | **R_max_** (RU) |  |
| *WT* | 2.53 (±0.13) | 9.90(±0.04) | 3.92(±0.18) | 23.35(±0.78) | 0.94(±0.01) |
| *D17N (FWL)* | 2.89(±0.01) | 9.78(±0.03) | 3.39(±0.01) | 27.75(±0.07) | 1.48(±0.07) |
| *D70N (FWL)* | 2.57(±0.13) | 0.102(±0.14) | 3.98(±0.26) | 20.9(±0.71) | 0.038(±0.04) |
| *K42E (FWL)* | 2.49(±0.04) | 9.54(±0.06) | 3.84(±0.08) | 19.65(±0.07) | 0.72(±0.01) |
| *V5Q*  *(FWH)* | 2.13(±0.01) | 9.82(±0.03) | 4.62(±0.03) | 24.55(±0.07) | 2.01(±0.08) |
| *W32Q (CDRH2)* | 28.6(±6.01) | 0.37(±0.05) | 0.01(±0.02) | 0.45(±0.07) | 0.06(±0.00) |
| *D28N (CDRL1)* | 2.60(±0.01) | 11.00 (±0.01) | 4.24(±0.06) | 26.1(±0.14) | 1.19(±0.07) |
| *D56N (CDRL2)* | 3.12(±0.01) | 10.5(±0.01) | 3.38(±0.11) | 29.1(±0.42) | 1.74(±0.04) |
| *R53G* (CDRL2) | 3.07(±0.01) | 11.5(±1.63) | 4.17(±0.16) | 13.5(±0.28) | 1.96(±0.04) |

***Differential Scanning Fluorimetry (DSF)***

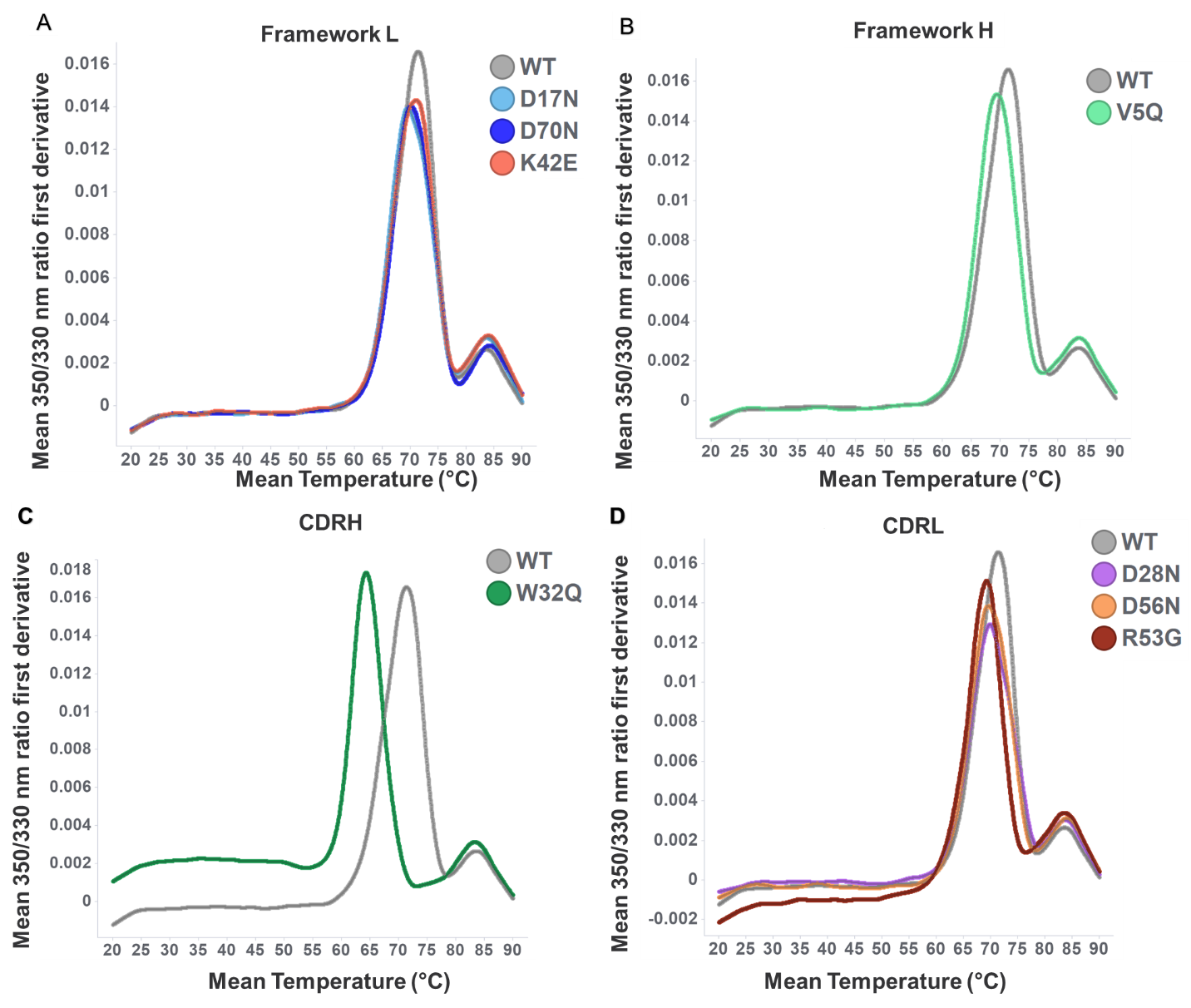

**Figure S5** Thermal unfolding profiles for all generated mutant molecules categorised by mutation site. 350nm/330nm ratio was obtained from differential scanning fluorimetry experiments for all expressed mutants and mAb1 WT at 150 mg/mL. To identify melting temperature peaks, the first derivative was calculated. The onset of unfolding (T_onset_) was identified at the inflection point of the first peak and was approximately 62°C for most mutants.

**Table S11** ***Thermal parameters derived from differential scanning fluorimetry of the WT and mutant mAb1 panel****. Unfolding onset temperatures (T_onset_) were comparable for all mAb1 mutants (~63°C), except for W32Q (CDRH) and R53G (CDRL), which had lower T_onsets_. Reduced thermal stability for W32Q was supported with reduced T_m1_ and T_agg_ values compared to WT. The distinction of lower thermal stability for R53G was weaker with a large deviation for T_agg_*. *All mAb1 samples were measured at 150 mg/mL. Only T_m1_ and T_m3_ peaks were detected for all molecules. FWL: light chain framework region; FWH: heavy chain framework region; CDRH2: heavy chain complementarity-determining region 2; CDRL1: light chain complementarity-determining region 1; CDRL2: light chain complementarity-determining region 2 (N=2).N=2 biological replicates.*

| Molecule | T_onset_ (°C) | T_m1_ (°C) | T_m3_ (°C) | T_agg_ (°C) |
| --- | --- | --- | --- | --- |
| *WT* | 62.85(±0.34) | 71.28(±0.17) | 83.71(±0.08) | 71.31(±2.40) |
| *D17N*  *(FWL)* | 63.19(±0.08) | 69.76(±0.13) | 83.61(±0.01) | 70.90(±0.53) |
| *D70N*  *(FWL)* | 63.32(±0.27) | 70.17(±0.04) | 84.25(±0.06) | 73.04(±4.12) |
| *K42E*  *(FWL)* | 63.18(±0.07) | 70.85(±0.11) | 83.97(±0.00) | 70.57(±1.14) |
| *V5Q*  *(FWH)* | 62.74(±0.28) | 69.42(±0.07) | 83.55(±0.06) | 69.59(±0.79) |
| *W32Q*  *(CDRH2)* | 60.08(±0.13) | 64.30(±0.01) | 83.27(±0.01 | 64.80(±0.96) |
| *D28N*  *(CDRL1)* | 62.56(±0.34) | 69.86(±0.26) | 83.65(±0.23) | 71.87(±1.59) |
| *D56N*  *(CDRL2)* | 62.51(±0.54) | 69.80(±0.81) | 83.88(±0.47) | 70.57(±2.39) |
| *R53G*  *(CDRL1)* | 58.63(±0.04) | 69.09(±0.20) | 83.52(±0.05) | 68.34(±10.46) |

***Dynamic light scattering (DLS)***

 
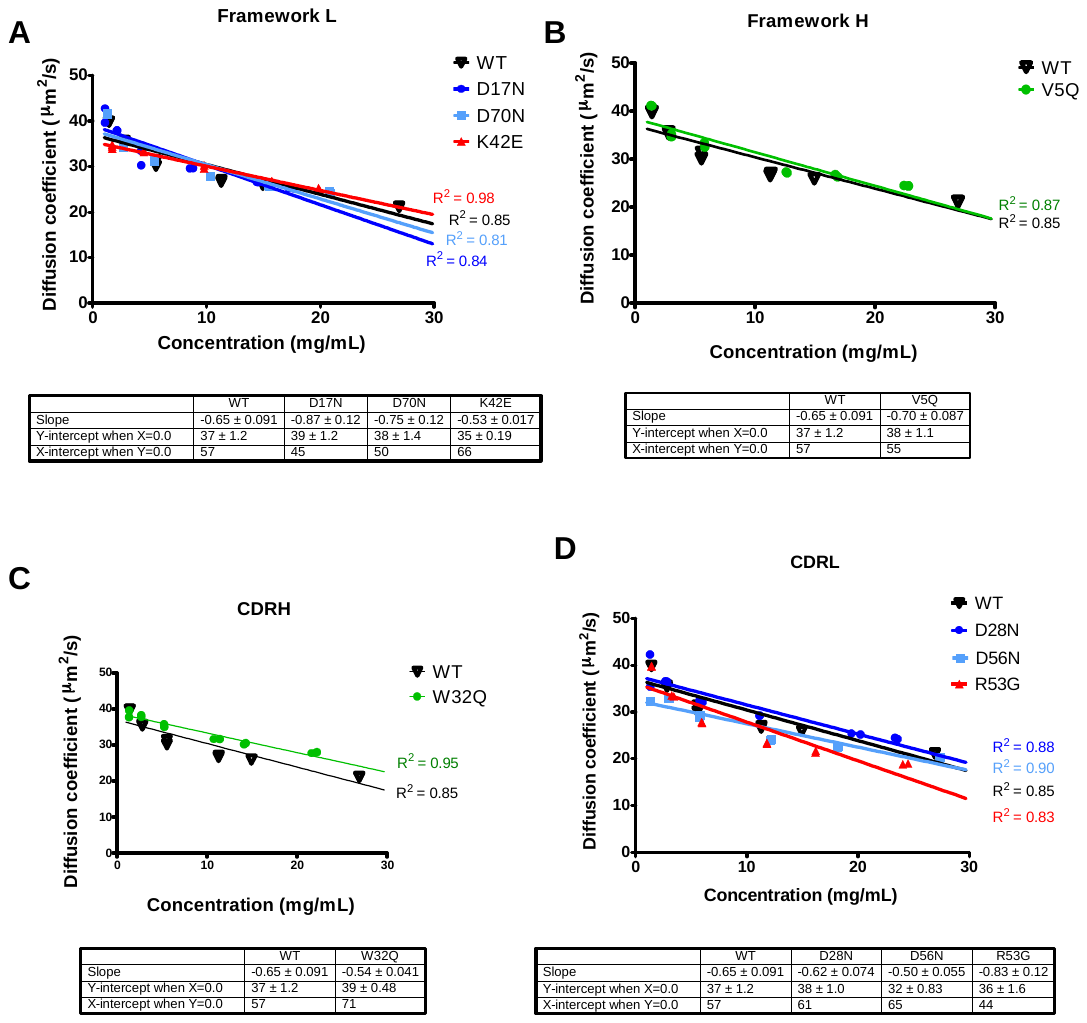

**Figure S6** Diffusion coefficients for each expressed mAb1 mutant and WT over a dilute concentration range (1-30 mg/mL), fitted with linear regression. Goodness of fit R-squared values are reported along with linear equations used to calculate self-interaction kD values.

1. Chemical Computing Group ULC (2021). "MOE 2020.09: Ensemble Protein Properties."
2. Salgado, J. C., et al. (2006). "Predicting the behaviour of proteins in hydrophobic interaction chromatography: 1: Using the hydrophobic imbalance (HI) to describe their surface amino acid distribution." Journal of Chromatography A 1107(1): 110-119.
3. Long, W. (Chemical Computing Group) (2022). "Bio-MOE: Custom MOE Biologics Applications
4. Kyte, J. and R. F. Doolittle (1982). "A simple method for displaying the hydropathic character of a protein." Journal of Molecular Biology 157(1): 105-132.
5. Sharma, V. K., et al. (2014). "In silico selection of therapeutic antibodies for develop ment: viscosity, clearance, and chemical stability." Proc Natl Acad Sci U S A 111(52): 18601-18606.
6. Tanford, C. (1962). "Physical chemistry of macromolecules. , John Wiley & Sons, Inc., New York 16, N. Y., 1961." 51(2): 190-190.
7. Sillero, A. and J. M. Ribeiro (1989). "Isoelectric points of proteins: theoretical determination." Anal Biochem 179(2): 319-325.
8. Ahmed, L., et al. (2021). "Intrinsic physicochemical profile of marketed antibody-based biotherapeutics." Proceedings of the National Academy of Sciences 118(37): e2020577118.
